# Modulation of Ether-à-go-go related gene (ERG) current governs intrinsic persistent activity in rodent neocortical pyramidal cells

**DOI:** 10.1101/156075

**Authors:** Edward D. Cui, Ben W. Strowbridge

## Abstract

While cholinergic receptor activation has long been known to dramatically enhance the excitability of cortical neurons, the cellular mechanisms responsible for this effect are not well understood. We used intracellular recordings in rat neocortical brain slices to assess the ionic mechanisms supporting persistent firing modes triggered by depolarizing stimuli following cholinergic receptor activation. We found multiple lines of evidence suggesting that a component of the underlying hyperexcitability associated with persistent firing reflects a reduction in the standing (leak) K^+^ current mediated by Ether-à-go-go-Related Gene (ERG) channels. Three chemically diverse ERG channel blockers (terfenadine, ErgToxin-1, and E-4031) abolished persistent firing and the underlying increase in input resistance in deep pyramidal cells in temporal and prefrontal association neocortex. Calcium accumulation during triggering stimuli appear to attenuate ERG currents, leading to membrane potential depolarization and increased input resistance, two critical elements generating persistent firing. Our results also suggest that ERG current normally governs cortical neuron responses to depolarizing stimuli by opposing prolonged discharges and by enhancing the post-stimulus repolarization. The broad expression of ERG channels and the ability of ERG blocks to abolish persistent firing evoked by both synaptic and intracellular step stimuli suggests modulation of ERG channels may underlie many forms of persistent activity observed in vivo.

## Introduction

Many brain regions contain neurons that generate long-lasting spiking responses to brief stimuli. In some of these areas, such as the brainstem nuclei that mediate the vestibulo–ocular reflex (VOR; 1), persistent responses play a central role in maintaining a sensory-motor system in a stable state. In cortical brain regions, persistent activity is associated with encoding of short-term memories (e.g., delay period firing during working memory tasks; 2) and in neuronal ensembles that represent time intervals (3). Recent work also demon-strates that persistent activity also occurs in hippocampal brain slices where mossy cells receive long-lasting synaptic barrages following brief stimuli (4, 5). Despite the widespread nature of persistent firing, the underlying mechanisms responsible for this firing mode have remained mysterious.

While synaptic reverberation has long been proposed to underlie persistent spiking responses recorded in vivo, especially in cortical regions (6, 7, 8), there is relatively little experimental support for this mechanism. By contrast, several decades of work using brain slices has demonstrated the ability of many cortical (9, 10, 11) and olfactory (12) neurons to generate persistent firing through cell-autonomous biophysical mechanisms. These intrinsic forms of persistent activity can be initiated by bursts of action potentials (APs; 13, 9) and are typically studied using brain slices exposed to muscarinic receptor agonists to enhance excitability (13, 9, 10, 14, 11) that likely mimic the increase in firing rate of cholinergic basal forebrain neurons during periods of heightened attention (15). Understanding the biophysical basis of persistent firing in brain slices would likely provide new tools that could be used to determine the relative roles of intrinsic and circuit mechanisms to persistent activity recorded in vivo.

While most experimental work on the underlying mechanism is in agreement that an increase in intracellular Ca^2+^ is required to trigger intrinsic persistent firing (e.g., 16, 17, 12), the critical downstream ion channels modulated by Ca^2+^ have not been identified. Many studies (16, 11, 18) have suggested that inward current mediated by Ca^2+^-activated non-selective cation channels (I_CAN_) underlies intrinsic persistent firing. Since the molecular basis of I_CAN_ has remained elusive, this hypothesis has been difficult to test definitively. Transient receptor potential (TRP) channels, a likely component of I_CAN_ (19, 14), have few selective antagonists and there have been no reports to date of TRP channel knockouts abolishing persistent activity though the ability of TRP proteins to form multimers (20) complicates testing this hypothesis.

In addition to I_CAN_-mediated Ca^2+^-sensitive currents, modulation of inward or outward currents active near rest have been suggested to underlie intrinsic persistent activity. Yamada-Hanff and Bean (21) demonstrated that the biophysical properties of subthreshold Na^+^ current could enable persistent firing. However, the absence of selective blockers of subthreshold Na^+^ channel activity had precluded directly testing this model. Alternatively, a decrease a “leak” K^+^ current could underlie the enhanced excitability associated with persistent firing. Presumably many muscarinic receptor agonists work though this mechanism to enhance overall excitability by inhibiting KCNQ channels that form the M current (22, 23). Similarly, Ether-à-go-go-related gene (ERG) channels can be modulated by second messenger cascades that converge on PKC (24). ERG is highly expressed in deep layers of neocortex (25, 26), among other regions, and is known to influence the excitability of a variety of neurons including midbrain dopaminergic neurons (27), brainstem neurons (28) and hippocampal CA1 pyramidal cells (29). Mutations in ERG have been linked to schizophrenia (30, 31) and many common antipsychotic medications are potent ERG blockers (32), suggesting that altered ERG function may underlie some types of cognitive dysfunction. Whether the increase in excitability that enables persistent responses to transient depolarizing stimuli is mediated by I_CAN_ currents, activation of subthreshold Na^+^ channels or attenuation of outward currents active near rest is not known.

In the present study, we used electrophysiological methods assess the mechanism responsible for intrinsic persistent activity in rodent neocortical slices. We find that ERG channel blockers abolish persistent firing in pyramidal cells in both temporal association neocortical as well as in prefrontal neurons. Blockade of ERG channels also greatly attenuates the increase in input resistance that underlies persistent firing, presumably reflecting a Ca^2+^-dependent attenuation in the leak ERG current. The robustness of ERG-mediated persistent firing and the widespread expression of ERG channels across diverse cortical regions suggest that modulation of ERG current may underlie many forms of persistent firing reported in vivo.

## Methods

### Slice preparation

Sprague-Dawley rats from postnatal 14-25 days were used in all experiments. Rats were anesthetized with ketamine and decapitated. The brain was then dissected and transferred into ice-cold (~0 °C) artificial cerebral spinal fluid (ACSF) composed of (in mM): 124 NaCl, 2.54 KCl, 1.23 NaH_2_PO_4_, 6.2 MgSO_4_, 26 NaHCO_3_, 10 dextrose, 1 CaCl_2_, equilibrated with 95% O_2_/ 5% CO_2_. Horizontal slices (300 *μ*m) were prepared from temporal association cortex (TeA; at the same dorsal-ventral level as the ventral hippocampus) using a Leica VT1200 vibratome. Recordings from prefrontal cortex (PFC) were performed on 300 *μ*m thick coronal slices that included the medial PFC. Slices were incubated at 30 ^o^C for approximately 30 min, and then maintained at room temperature (~25 ^o^C) until use. All experiments were carried out under guidelines approved by the Case Western Reserve University Animal Care and Use Committee.

### Electrophysiology

Intracellular and cell-attached recordings were performed in a submerged chamber maintained at 30 ^o^C and perfused continuously (~2 ml/min) with ACSF containing (in mM): 124 NaCl, 3 KCl, 1.23 NaH_2_PO_4_, 1.2 MgSO_4_, 26 NaHCO_3_, 10 dextrose, 2.5 CaCl_2_, equilibrated with 95% O_2_/ 5% CO_2_. Whole-cell and cell-attached recordings were made using an Axopatch 1D amplifier (Axon Instruments/Molecular Dynamics). Patch clamp recording electrodes with resistances 3-8 MΩ were pulled from 1.5 mm OD thin wall borosilicate glass tubing (World Precision Instruments; WPI), using a micropipette puller (P-97; Sutter Instruments). The pipettes contained (in mM): 140 K-methylsulfate (MP Biochemicals), 4 NaCl, 10 HEPES, 0.2 EGTA, 4 MgATP, 0.3 Na_3_GTP, 10 phosphocreatine. In some experiments, 10 mM BAPTA (1,2-bis(o-aminophenoxy)ethane-N,N,N’,N’-tetraacetic acid) was substituted for EGTA in the internal solution to enhance Ca^2+^ buffering. In one set of experiments, 10 *μ*M E-4031 was added to the internal solution. Individual neurons were visualized under infrared differential interference contrast (IR-DIC) video microscopy (Zeiss Axioskop FS1). Recordings were low-pass filtered at 5 kHz (FLA-01, Cygus Technology) and acquired at 10 kHz using a simultaneously-sampling 16-bit data acquisition system (ITC-18, Instrutech) operated by custom software written in VB.NET on a Windows-based PC. Membrane potentials shown are not corrected for the liquid junction potential. Synaptic stimulation was performed using an insulated tungsten monopolar electrode placed in layer 3 near and connected to a constant current stimulus isolation unit (A360, WPI).

In this study, we recorded exclusively from layer 5 (L5) neocortical pyramidal cells that generated “regular spiking” responses to 2 s duration depolarizing currents as defined by previous neocortical studies (33, 34, 35). Cells were selected using IR-DIC visualization and were confirmed to be pyramidal cells using 2-photon imaging in a subset of recordings. The average resting membrane potential of TeA L5 neurons was -66.1 ± 0.25 (N = 149; range -60 to -70 mV). The average R_In_ of these neurons was 107 ± 3.7 MΩ. Recordings from neurons with resting membrane potentials > -60 mV or R_In_ < 60 MΩ were excluded. Persistent firing was evoked in most experiments in this study using a standardized protocol described in (36; 2 s duration depolarizing step that generated ~10 Hz continuous firing from -70 mV).

Except for experiments using intracellularly-applied E-4031, all drugs were applied by switching the bath perfusion reservoir. Unless specified, all drugs were purchased from Tocris Bioscience. Drugs used included: carbamoylcholine chloride (Carbachol, CCh), used at 2 *μ*M with 10 mM stock solutions of CCh in water were prepared each day; gabazine (SR 95531, used at 10 *μ*M); 2,3-Dioxo-6-nitro-1,2,3,4-tetrahydrobenzo[f]quinoxaline-7-sulfonamide (NBQX, used at 10 *μ*M); 4-aminopyridine (4-AP, used at 100 *μ*M and purchased from Sigma); tetraethylammonium (TEA, used at 1 mM and purchased from Sigma); ZD7288 (used at 10 *μ*M); D-APV (used at 10 *μ*M), N-(3,4-Difluorophenyl)-N’-(3-methyl-1-phenyl-1H-pyrazol-5-yl)urea (ML297 used at 0.67 *μ*M); terfenadine (Terf, used at 10 *μ*M); E-4031 dihydrochloride (used at 10 *μ*M); fexofenadine (used 30 *μ*M); linopirdine dihydrochloride (used at 30 *μ*M); pirenzepine dihydrochloride (used at 10 *μ*M); ErgToxin-1 (used at 50 nM) was purchased from Alomone Lab and dissolved in ACSF.

### Experimental design and statistical analysis

We estimated input resistance under both static and time-varying conditions. For static tests, performed when we were able to hold the membrane at a constant potential for extended periods, we estimated R_In_ from the maximal voltage deflection elicited by a standard (typically -50 pA for 2 s) injected current step. To estimate R_In_ changes underlying persistent activity (e.g., Fig. 5B-C), we hyperpolarized the membrane potential 500 ms following the offset of depolarizing step to abolish persistent firing and then injected a series of small hyperpolarizing test pulses (-50 pA, 300 ms duration) to assay input resistance every 600 ms. By delaying the R_In_ assay procedure by 500 ms, we could ensure that the depolarizing step was effective in triggering persistent activity since in all experiments the first post-step AP was elicited within 500 ms. We then used a two-stage correction process to account for the effect of the changing membrane potential during the afterdepolarization (ADP) and the voltage dependence of input resistance in neocortical pyramidal cells (37). First we detrended each response to the 300 ms hyperpolarizing pulses by subtracting the regression line fit between the mean potential during 10 ms periods acquired immediately before and 290 ms following each step. We then calculated a raw input resistance estimate from the voltage deflection elicited by each step calculated from the mean potential during the final 50 ms of the detrended step response. Finally, we compared this R_In_ estimate to calibration R_In_ measurements acquired in interleaved trials where R_In_ was measured under static conditions. We used linear regression fits of this calibration data to determined the expected input resistance at any arbitrary voltage within the calibrated range (-75 to -60 mV; R^2^ = 0.76 ± 0.029 for R_In_ estimates in ACSF and 0.83 ± 0.063 in CCh). Our final “corrected” delta input resistance measurement reflect the difference between the measured R_In_ and the expected R_In_ at that particular membrane potential. For example, in one experiment the static input resistance recorded at -70 mV was 100 MΩ which increased to 110 MΩ at a -65 mV holding potential. If the peak ADP response occurred at -65 mV in that cell, generating a detrended R_In_ estimate of 150 MΩ, we would then report a “corrected” change in input resistance of + 40 MΩ (150 - 110 MΩ) at that time point. We repeated this procedure using at least three ADP trials in each cell, and at each time point, and report the mean corrected delta R_In_ values. Summary time plots show the mean ± S.E.M. across multiple experiments of these single-cell corrected delta R_In_ estimates.

We calculated the probability of triggering persistent firing by examining 3-5 consecutive responses to standard 2 s duration depolarizing conditioning steps (1-2 min between trials; persistent firing was terminated by manually hyperpolarizing the bias current–moving the membrane potential to < -80 mV and then gradually reducing the bias current to restore the standard -70 mV holding potential). Except for experiments designed to measure dynamic changes in R_In_, we determined if the neuron continued to fire for at least 5 s following the offset of the depolarizing conditioning step to assess whether persistent firing occurred. Summary persistent firing probabilities presented in the figures (e.g., Fig. 2E) reflect the average of the probabilities computed from each neuron tested. Membrane potential sag ratios were assessed from responses to 2 s duration current steps that evoked ~20 mV hyperpolarization from a -60 mV holding potential. Sag ratio was computed by dividing the estimated R_In_ at the end of the step from the R_In_ estimated 50 ms following step onset. Action potential threshold was calculated as the voltage at which dV/dt was greater than 10% of maximum dV/dt during the rising phase of the AP. The AP half-width was calculated from the duration at the voltage halfway between AP threshold and AP peak. We the calculated I/V relationships shown in Fig. 7 using current-clamp recordings by determining the injected current required to reach each membrane potential during ramp stimuli acquired in control and experimental conditions. Data presented represent the difference in estimated whole-cell current between the experimental and control conditions at each voltage. Data were expressed as mean ± S.E.M. Significance level with p < 0.05 was used. Multiple comparisons were Bonferroni corrected. Data analysis employed custom programs written in Python 3.6 and Matlab 2015b. Statistical tests were performed in Python and R.

## Results

As reported by others (e.g., 11 in neocortical neurons and 16 in entorhinal cortical neurons), activation of muscarinic receptors attenuated the hyperpolarizing afterpotential that normally follows burst of APs and revealed an afterdepolarization (ADP) that can trigger persistent firing (Fig. 1A). Throughout this study, we refer to the depolarizing current injection that initiates the persistent firing as the “conditioning” step to differentiate it from other current stimuli used to measure input resistance and to assay intrinsic excitability in experiments described below. In addition to enabling persistent firing, the cholinergic receptor agonist carbochol (CCh; 2 *μ*M) depolarized pyramidal cells and triggered spontaneous firing by itself in most cells within 20-30 min. To compensate for this second effect of CCh, we included a low concentration (0.67 *μ*M) of the GIRK channel activator ML297 in a subset of experiments. (Drugs used in each experiment are specified in the related figure legend.) At this concentration, the tonic hyperpolarization produced by ML297 compensated for the depolarizing effect of CCh, leading to relatively stable membrane potential while retaining the ability of CCh to enable persistent firing in response to depolarizing current steps over multiple trials (e.g. Fig. 1A). As reported by others (16, 38), the ability of cholinergic receptor agonists to facilitate persistent firing responses to depolarizing stimuli was largely mediated by actions of the m1 subclass of muscarinic receptors. In our experiments, the m1 receptor antagonist pirenizpine abolished persistent firing responses (Fig. 1B-C). The effectiveness of standard depolarizing step responses (2 s duration triggering firing at ~8-12 Hz from -70 mV) was similar in CCh and CCh + ML297 (Fig. 1C). Persistent firing responses did not occur in control ACSF solution (0/5 experiments interleaved with CCh experiments where persistent firing was reliably evoked) and never occurred when ML297 was presented without CCh (0/8 cells).

**Figure 1:**
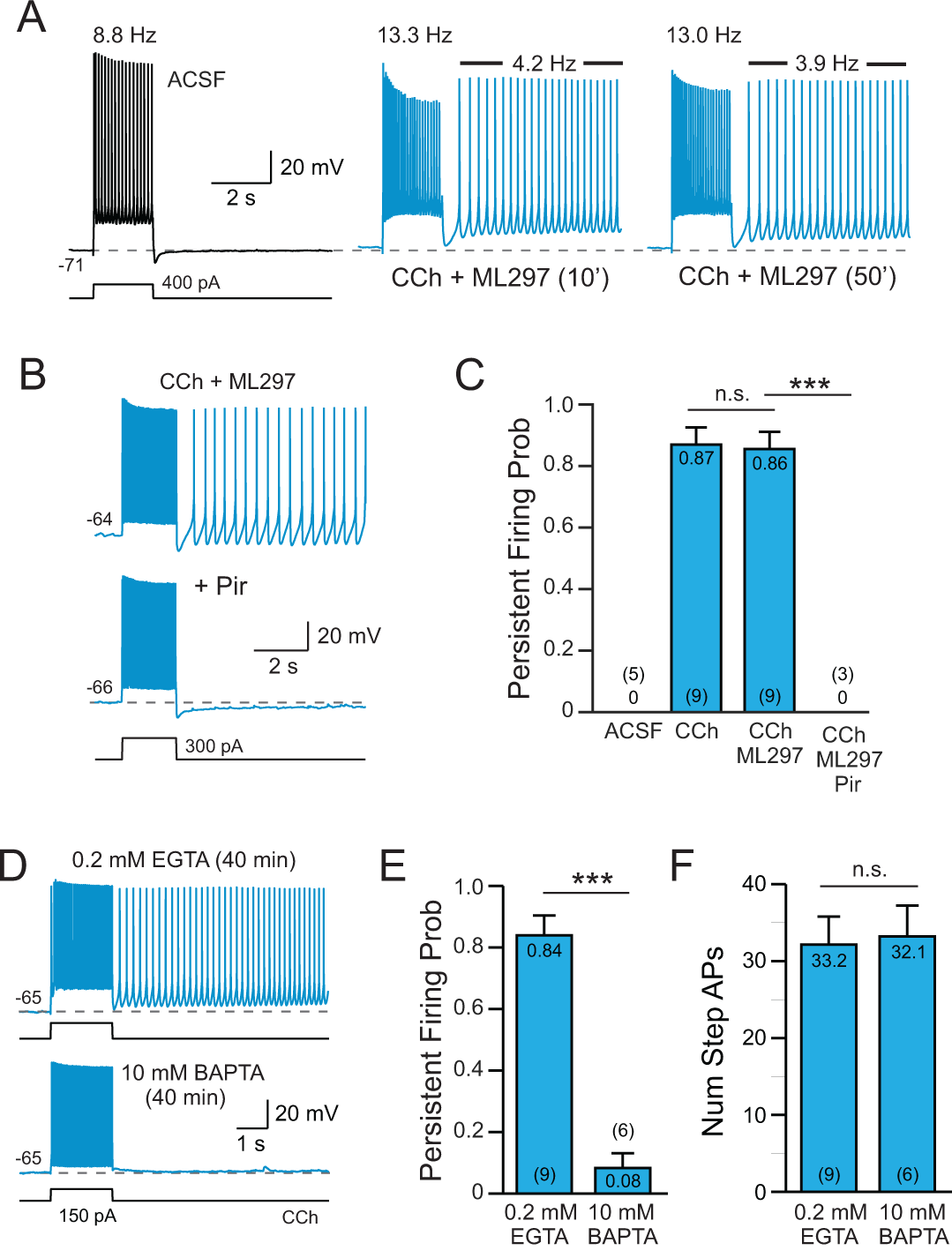
Cholinergic receptor activation enables persistent firing following depolarizing stimuli. A, Bath application of 2 *μ*M carbachol (CCh; blue traces) reveals persistent spiking following depolarizing steps in L5 pyramidal cells from rat temporal association (TeA) neocortex. Firing rates during the step response and during the post-step persistent spiking period were stable > 45 min (right panel acquired following 50 min exposure to CCh; firing rates indicated above traces). The GIRK activator ML297 (0.67 *μ*M) was included in the bath solution to compensate for the tendency of CCh to depolarize pyramidal cells (see Methods for details). B, Blockade of persistent firing by 10 *μ*M pirenzipine (Pir). C, Summary of the probability of triggering persistent firing in ACSF, CCh, CCh + ML297 and in Pir (and CCh + ML297). *** P = 6.49E-05, T = 7.62, df = 9, two-sample t-test; n.s. P = 1.00, T = 0.0879, df = 15, two-sample t-test. Group means and Ns indicated inside each bar. D, Persistent firing in response to depolarizing step stimuli could be evoked using an internal solution containing 0.2 mM EGTA (top trace) but not with an internal solution containing 10 mM BAPTA (bottom trace). E, Summary plot of the probability of triggering persistent firing in 0.2 mM EGTA and 10 mM BAPTA. *** P = 2.36E-06, T = 7.96, df = 13, two sample t-test. F, Plot of the number of APs evoked by the depolarizing step in EGTA- and BAPTA-based internal solution. n.s. P = 0.862, T = -0.178, df = 13, two-sample t-test.

Persistent firing in L5 neocortical neurons following cholinergic stimulation required an increase in intracellular calcium. We rarely observed persistent firing following depolarizing test stimuli when the intracellular Ca^2+^ concentration was strongly buffered using 10 mM BAPTA (8% of experiments; Fig. 1D-F). In parallel experiments in which the time since breakthrough to whole-cell recording mode was similar (40 minutes), we were able to evoke persistent firing in > 80% experiments using our standard internal solution that contained 0.2 mM EGTA (Fig. 1D). The spiking response to the conditioning depolarizing step was similar in both recording conditions (Fig. 1F). These results suggests that persistent firing is triggered by an increase in intracellular Ca^2+^ and is consistent with previous work testing Ca^2+^ chelators in neocortical neurons (11). Persistent firing also reflected cell-autonomous processes since this response could be evoked following blockade of ionotropic glutamate and GABA receptors with NBQX (10 *μ*M), d-APV (25 *μ*M) and gabazine (10 *μ*M; N = 3 experiments; data not shown).

### ERG channel blockers abolish persistent firing

While previous reports (21, 39, 11, 40) have suggested multiple potential biophysical channels that could contribute to the increased excitability responsible for persistent firing in a variety of cortical pyramidal cells, the underlying mechanism has not been clearly demonstrated. A common explanation for increased excitability associated with an increase in input resistance is a reduction in a subpopulation of K^+^ channels that are open near the resting membrane potential (41, 42). As part of a survey of potential leak K^+^ channel mechanisms, we found that three chemically diverse Ether-à-go-go-Related Gene (ERG; comprising Kv11/KCNH2 channels; 31, 43) channel antagonists abolished persistent firing in L5 neocortical pyramidal cells. ERG currents help set the resting membrane potential in many CNS cells (29, 27, 44, 28) and ERG1 and ERG3 subunits are expressed at high levels in neocortical neurons (26).

Terfenadine (Terf; 10 *μ*M), histamine H1 receptor antagonist that also commonly is used to attenuate ERG currents (45, 32) eliminated persistent firing evoked by depolarizing step stimuli (5/5 experiments in CCh + ML297 and 6/6 in CCh alone; Fig. 2A; tested at -70 mV in all experiments). This effect of Terf appeared to be independent of its action on histamine receptors since another H1 receptor-specific antagonist that is structurally similar to Terf (fexofenadine; 46) had no effect on persistent firing even when tested at 3 times higher concentration (30 *μ*M; N = 3 experiments). Terfenadine also did not appear to abolish persistent firing by attenuating the depolarizing stimulus since there was no significant reduction in the number of spikes evoked by the depolarizing step in CCh (22.8 ± 1.4 vs 23.0 ± 1.4 spikes in Terf; P = 0.945; paired t-test; N = 10 experiments).

**Figure 2:**
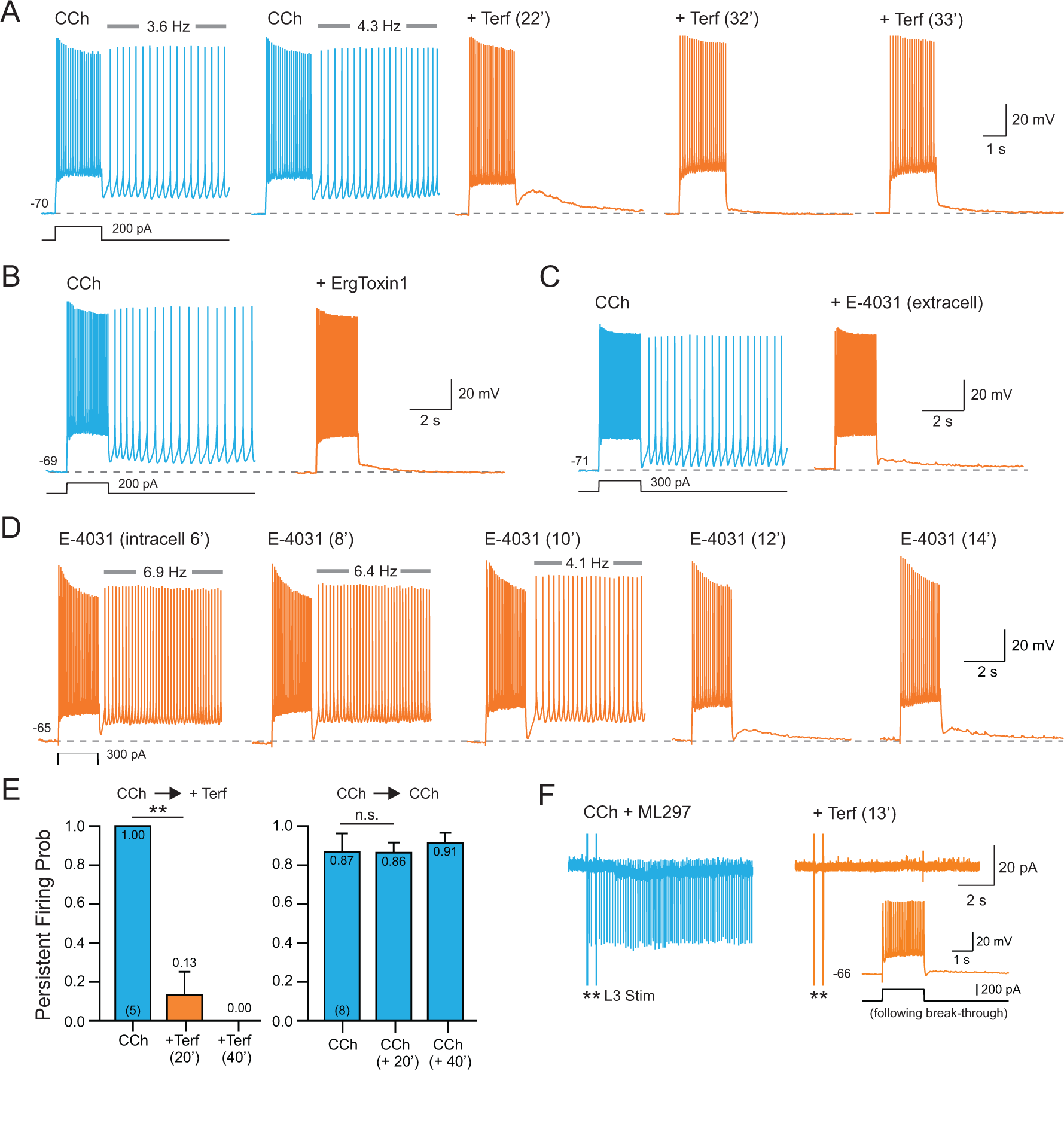
ERG blockers abolish persistent firing in neocortical neurons. A, Terfenadine (Terf; 10 *μ*M; orange traces) abolished persistent firing evoked by depolarizing steps in CCh + ML297. Terfenadine exposure time indicated above each trace. B, The peptide ERG channel blocker ErgToxin1 (50 nM) also abolished persistent firing. C, Extracellular application of E-4031 (10 *μ*M) blocked persistent firing recorded under the same conditions as A-B. D, Intracellular perfusion with E-4031 (10 *μ*M) abolished persistent firing within 12 min. Persistent firing continued to be evoked by test stimuli in interleaved control experiments without intracellular E-4031 for more than 40 min (N = 6). E, Left, plot of the probability of evoking persistent firing > 10 s before and after exposure to terfanadine. At 20 min: ** P = 0.0029, T = 6.50, df = 4, paired t-test; at 40 min, P = 0.0079, Fisher’s Exact Test. Right, persistent activity could be stably evoked in parallel experiments extending through the same duration without Terf (0 min vs. 20 min: P = 0.44; 20 min vs. 40 min: P = 0.98; 0 min vs. 40 min: P = 0.39, Tukey Honest test for multiple comparison). F, Terfenadine abolished persistent firing assayed in a cell-attached recording from a L5 neocortical neuron. Response triggered by two extracellular stimuli in L3 (asterisks). Terfenadine abolished the synaptically-triggered persistent firing (orange trace). Intracellular recordings from the same neuron following break-through to whole-cell mode demonstrated physiological normal step responses after synaptically-evoked persistent firing was abolished (inset).

While Terf required at least 20 min to block persistent firing in our experiments, this time course is consistent with the intracellular binding site on ERG channels for this antagonist (47, 48). ErgToxin1, by contrast, blocked persistent firing in less than 20 min (Fig. 2B; 4/4 experiments). This ERG-specific peptide toxin produced endogenously by scorpions binds to extracellular sites on ERG channel subunits (49, 50, 51, 52), accounting for the more rapid time course of blockade. We also tested a third ERG channel antagonist, E4031, which binds to an intracellular domain on ERG channel subunits (48, 47, 28). This compound was effective when applied both in the extracellular bathing media (Fig. 2C, 5/5 experiments) and rapidly blocked persistent firing when we recorded from L5 pyramidal cells using an internal solution containing 10 *μ*M E-4031 (Fig. 2D, 5/5 experiments). The ability of ERG channel blockers to abolish persistent firing was unlikely to be attributable to run-down in these experiments because Terf abolished persistent firing within the 20-30 min time frame when persistent firing could be reliably evoked in interleaved control experiments (Fig. 2E). Terfenadine also abolished persistent firing evoked by extracellular synaptic stimulation and assayed through cell-attached recordings from L5 pyramidal cells (Fig. 2F), demonstrating that ERG channel antagonists were effective under relatively physiological conditions that that did not involve a whole-cell recording configuration.

### ERG-mediated responses in L5 pyramidal cells

The ability of three different ERG channel blockers to abolish persistent firing suggests this subtype of K^+^ channel may play a critical role in regulating excitability in pyramidal cells. To our knowledge, there are no published reports addressing properties of ERG currents in neocortical neurons. We tested the role of ERG currents in these neurons by employing a set of previously-established tests of ERG channel function (28, 27, 53). First, we found that blockade of ERG channels with Terf increased the number of spikes evokes by a standard depolarizing step when tested in control ACSF (Fig. 3A). Over 14 similar experiments, attenuation of ERG currents with Terf increased the number of spikes evoked by 2 s depolarizing steps by 23.5% (Fig. 3B). The increased excitability generated by Terf likely reflected a reduction in the resting (leak) K^+^ conductance mediated by ERG channels since Terf increased the apparent input resistance by 21.1% in L5 pyramidal cells maintained at -70 mV (via adjusting the bias current injected; Fig. 3C). In separate experiments where the membrane potential was not manually clamped, Terf depolarized L5 pyramidal cells by ~10 mV (Fig. 3D), consistent with a reduction in a tonically open ERG channels. Activation of muscarinic receptors did not occlude the ability of Terf to attenuate a component of the leak K^+^ current since Terf was still able to depolarize pyramidal cells (Fig. 3E) and increase apparent input resistance (Fig. 3F) even in the presence of 2 *μ*M CCh. With cholinergic receptor stimulation via CCh, Terf was able to depolarize L5 pyramidal cells sufficiently to induce spontaneous firing (orange trace in Fig. 3E), demonstrating that a reduction in the steady-state ERG current is sufficient to induce persistent firing.

**Figure 3:**
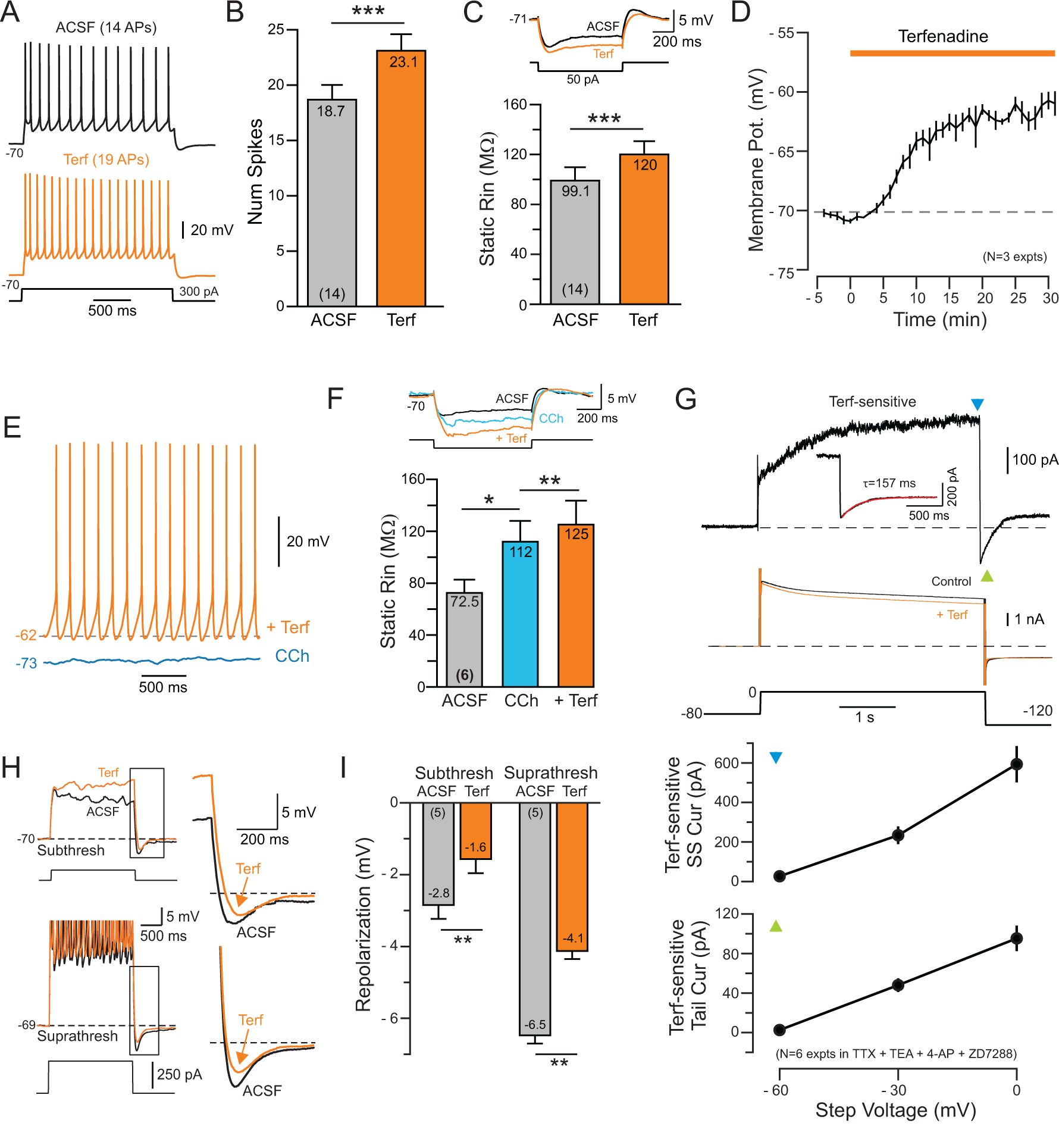
ERG-mediated currents in neocortical pyramidal cells. A, Terfenadine increases the number of spikes evoked by depolarizing current stimuli (comparison from ACSF to terfenadine). B, Plot of results from 14 experiments similar to A. *** P = 2.93E-04, T = 4.89, df = 13, paired t-test. C, Terfenadine increases input resistance assayed from similar reference membrane potentials (~-70 mV). *** P = 1.50E-04, T = 5.28, df = 13, paired t-test. D, Plot of mean membrane depolarization evoked by Terf in 3 pyramidal cells. E, Following cholinergic stimulation with CCh (blue trace), the same concentration of Terf elicited spontaneous firing (orange trace). F, Plot of input resistance assayed at -70 mV in ACSF, following CCh and with CCh + Terf. * P = 0.027, T = 3.74, df = 5; ** P = 0.0058, T = 5.4253, df = 5, paired t-test. G, Isolation of Terf-sensitive current in responses to steps to 0 mV. Raw responses (not leak subtracted) shown in middle panel; top trace represents subtraction of Terf response from control response. Responses acquired in 1 *μ*M TTX, 100 *μ*M 4-AP, 1 mM TEA and 10 *μ*M ZD7288 and with external K^+^ increased from 3 to 20 mM. Summary plots show voltage dependence of steady-state outward current analyzed at the end of the step (top plot, at time marked by green downward triangle on trace) and the peak tail current following the depolarizing step (bottom plot, at time marked by gray upwards triangle). H, Terfenadine attenuates post-step repolarization to both sub- (top) and supra-threshold (bottom) responses to depolarizing steps. Enlargements of repolarization shown on right. I, Plot of the attenuation of the post-step repolarization. Subthreshold: ** P = 0.00787, T = 4.93, df = 4; ** Suprathreshold: P = 0.00167, T = 7.53, df = 4. Both paired t-tests.

Using previously-established voltage-clamp recording protocols for assaying ERG currents (53, 28, 44), Terf selectively attenuated a late developing, voltage-sensitive outward current (Fig. 3G). In response to strong depolarizing steps (from -80 to 0 mV) the Terf-sensitive current developed with a time constant of 810 ms, similar to estimates reported in heterologously-expressed ERG channels (e.g., Figure 2A of 54, Figure 1A of 55, and Figure 1A of 24) and about 4-fold slower than the activation kinetics of I_M_ (56). The Terf-sensitive current response deactivated with time constant of ~160 ms (Fig. 3G inset; mean 153.5 ± 10.0 ms; N = 6) which is also consistent with previous estimates of ERG channel kinetics recorded in heterologous expression systems (57, 58) and in whole-cell recordings from brain slices (28). Both the peak steady-state Terf-sensitive outward current and the maximal tail current amplitude increased with larger amplitude depolarizing steps (bottom two plots in Fig. 3G), as expected for ERG currents.

ERG current functions to help repolarize APs in cardiac cells, where Ether-à-go-gorelated gene channels are most frequently studied (51). ERG currents appear to play a similar role in neocortical neurons since Terf reduced the membrane potential repolarization following both subthreshold (Fig. 3H, top) and suprathreshold depolarizing steps (Fig. 3H, bottom). In 5 pyramidal cells tested, Terf eliminated approximately one-third of the repolarization following both types of steps (Fig. 3I, repolarization measured relative to the pre-step membrane potential). These results suggest that neocortical pyramidal cells express ERG channels that are tonically active near the normal resting potential which function both to dampen excitability in response to depolarizing stimuli and to enhance the post-stimulus repolarization.

Neocortical pyramidal cells express a wide variety of K^+^ channels which also could affect excitability following depolarizing conditioning steps in CCh. I_H_ often triggers rebound hyperexcitability (59) and is expressed in L5 pyramidal cells (Fig. 4A-C and see 60). I_H_ therefore could potentially contribute to persistent activity. However, the selective I_H_ blocker ZD7288 failed to abolish persistent firing in 6/6 cells tested (Fig. 4B) while consistently attenuating the membrane potential “sag” in responses to hyperpolarizing steps mediated by I_H_ (Fig. 4C). In a separate set of experiments, we were able to evoke typical persistent firing responses from our standard depolarizing test responses in slices pretreated with ZD7288 (Fig. 4D) or in the nonselective I_H_ blocker Cs^+^ (10 mM; N = 3; data not shown), suggesting that modulation of IH was not responsible for persistent firing in L5 neurons. Even with IH blocked with ZD7288, Terf was still able to abolish persistent firing (Fig. 4D; N = 4). The selective I_M_ blocker linopirdine (30 *μ*M; 23) also failed to abolish persistent firing in 3/3 cells tested (Fig. 4E). The dihydropyridine Ca^2+^ channel antagonist nimodipine also did not prevent persistent firing (N = 6; 20 *μ*M), suggesting that persistent firing did not result from an interaction between K^+^ and L-type Ca^2+^ channels.

**Figure 4:**
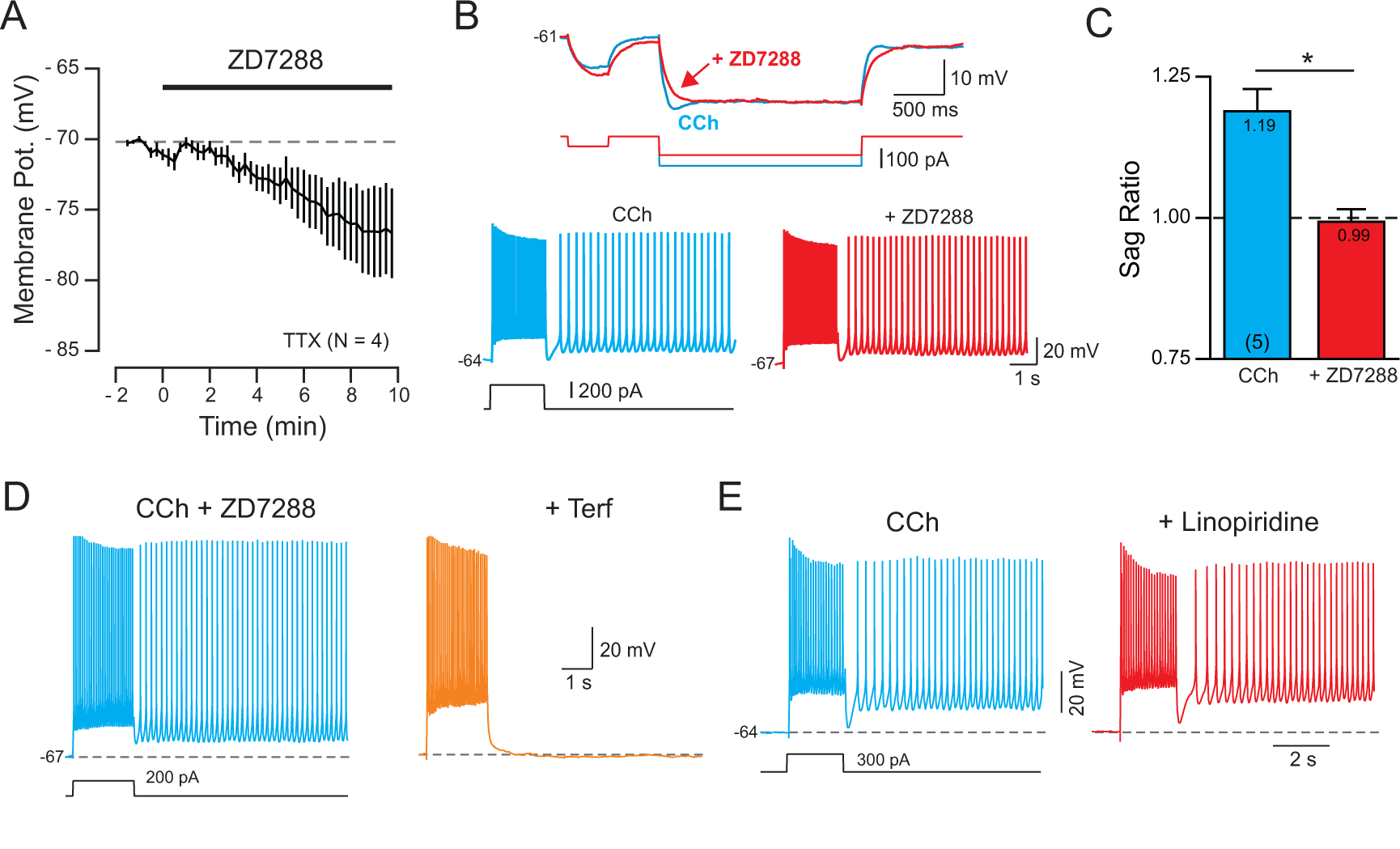
I_H_ and I_M_ currents are not required for persistent firing. A, Bath application of the specific I_H_ blocker ZD7288 (10 *μ*M) hyperpolarized L5 pyramidal cells. B, Blockade of I_H_ by ZD7288 eliminated the membrane potential sag evident in responses to hyperpolarizing current steps (top) but did not abolish persistent firing evoked by a depolarizing step (bottom, both representative of 5/5 cells tested). C, ZD7288 significantly reduced sag ratio. * P = 0.0257, T = 3.464, df = 4, paired t-test; see Methods for details of sag ratio analysis). ZD7288 did not block persistent firing in 5/5 cells tested (n.s. P = 1.00, Fisher’s Exact Test). D, Blockade of I_H_ by ZD7288 did not occlude the ability of Terf to abolish persistent firing (representative of 4/4 experiments). E, Attenuation of I_M_ current with linopiridine (30 *μ*M) failed to abolish persistent firing (typical of 3 experiments). In the same experiments, linopiridine increased the number of spikes evoked by depolarizing step from 23.50 ± 4.76 to 26.68 ± 4.97 (P = 0.0197, T = 7.02, df = 2, paired t-test).

### Transient increase in input resistance associated with persistent activity

The results presented thus far suggest that the hyperexcitability underlying persistent activity could reflect a transient attenuation in the component of the leak K^+^ current mediated by ERG channels. We next tested this hypothesis by assaying the change in intrinsic properties immediately following depolarizing step stimuli. When assayed using brief depolarizing steps, the depolarizing conditioning step converted just subthreshold responses into suprathreshold responses (Fig. 5A), consistent with either an increase in inward current (such as I_CAN_) or a reduction in a leak K^+^ current like ERG. We assayed input resistance using trains of brief (300 ms) hyperpolarizing current pulses to discriminate between these possibilities. In these experiments, we injected a steady hyperpolarizing current starting 0.5 s after the offset of the conditioning depolarizing step to prevent continuous persistent firing; the train of hyperpolarizing test pulses was applied on top of this steady hyperpolarization (Fig. 5B). By delaying the steady hyperpolarization by 0.5 s, we could verify that the conditioning step stimulus was effective in initiating persistent firing (see example trace in Fig. 5B with a single spike following the conditioning step).

**Figure 5:**
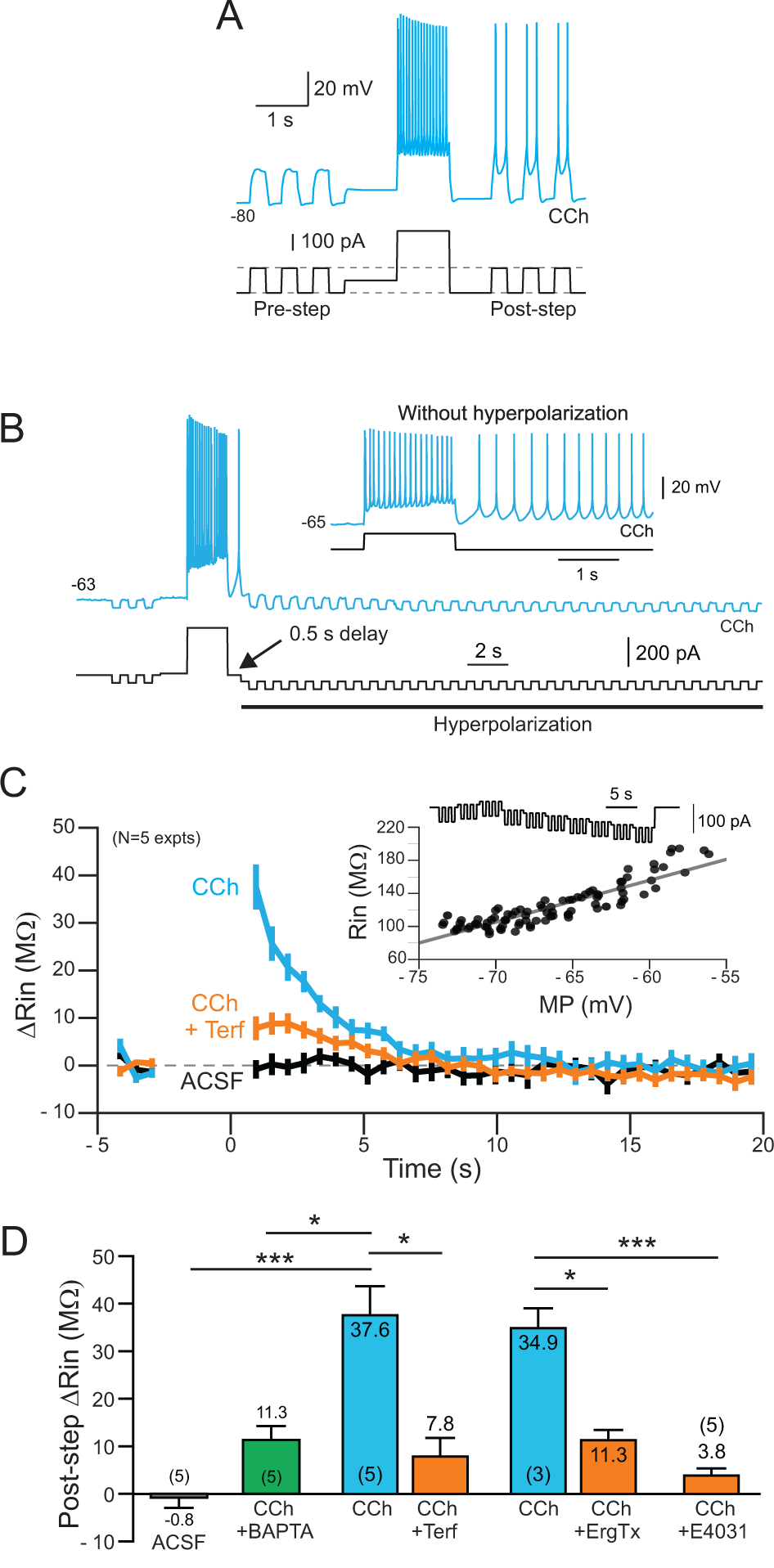
Time course of ERG-mediated change in input resistance. A, Demonstration of increased excitability following a depolarizing conditioning step in CCh. Brief test pulses that were subthreshold before the conditioning step become suprathreshold following the step. Neuron maintained at -80 mV to prevent persistent firing between test pulses. B, Example response to train of hyperpolarizing test pulses used to assay input resistance. Additional continuous hyperpolarizing bias current was applied 500 ms following the offset of the depolarizing conditioning step. Without this bias current, the neuron fired persistently in response to the conditioning step (inset). C, Plot of change in input resistance assayed by hyperpolarizing pulses in ACSF (black plot), CCh (blue) and CCh+Terf (orange). See Methods for details. Inset shows example input resistance calibration response. D, Summary of change input resistance following the depolarizing conditioning step. *** (ACSF/CCh) P = 1.29E-04, T = 6.23, df = 11, two-sample t-test; * (CCh/BAPTA vs CCh) P = 0.0171, T = 3.46, df = 8, two sample t-test; * (CCh/CCh+Terf) P = 0.022, T = 4.49, df = 4, paired t-test; * (CCh/CCh+ErgTx) P = 0.029, T = 5.79, df = 2, paired t-test; *** CCh vs. CCh/E4031: P = 3.29E-04 T = 5.58, df = 11, two-sample t-test. Two independent CCh data sets (N = 3 and 5) were combined when computing two-sample t-test statistics.

Simply assaying R_In_ based on the membrane potential change elicited by hyperpolarizing current pulses following the conditioning step suggested that the input resistance increased by ~34 MΩ during the ADP (from 130 ± 25.6 to 164 ± 30.3 MΩ; P = 4×10^−4^, T = 8.32, df = 5; paired t-test; N = 6). This estimate, however, is subject to several potential artifacts. First, the membrane potential is continuously hyperpolarizing following the conditioning step, especially during first few R_In_ test pulses. We corrected for this effect by detrending the membrane potential before calculating the voltage deflection elicited by each test pulse (see Methods for details). The second complication is that input resistance in L5 pyramidal cells varies depending on the membrane potential, with higher R_In_ estimates at more depolarized membrane potentials. To compensate for this effect, we determined the R_In_/V_M_ relationship over the relevant voltage range in each neuron (and in each drug condition; Fig. 5C, inset) and present our results as changes in R_In_ following the conditioning step relative to the steady-state R_In_ measured at that particular membrane potential prior to the conditioning step. Following these two correction procedures, we still find a large (35-38 MΩ) increase in input resistance following the conditioning step that decays with a time constant of 4.3 s (blue trace in Fig. 5C). (The similarity between R_In_ estimates with and without correction procedures reflects the opposing effects of the two types of artifacts when using hyperpolarizing test pulses.) During the same time period, we find no change in apparent R_In_ following conditioning depolarizing step responses in control ACSF (black trace in Fig. 5C).

In CCh, the ERG channel blocker Terf attenuated most of the increase in input resistance following the conditioning step (orange trace in Fig. 5C). We found similar results in 3 experiments using ErgToxin1 where most of increase in R_In_ was eliminated after ERG channels were blocked. Figure 5D summarizes the reduction in elevation in apparent R_In_ in separate sets of experiments with Terf and ErgToxin1. We also repeated the same procedure in 5 neurons recorded with the intracellular ERG channel blocker E-4031 added to the internal solution and observed only a small (~4 MΩ) increase in apparent Rin. The modest residual increase in apparent R_In_ following ERG channel blockade was similar (or less than) the peak R_In_ increase observed when intracellular Ca^2+^ was strongly buffered with a BAPTA-containing internal solution (green bar in Fig. 5D).

Neocortical pyramidal cells express large subthreshold Na^+^ currents that can contribute to persistent firing (21, 61) and can influence R_In_ estimates. To determine if the effects of ERG blockers were independent of voltage-gated Na^+^ channels, we applied tetrodotoxin (TTX; 1 *μ*M) along with 4-AP and CCh to increase excitability. In this drug combination, depolarizing conditioning steps reliably triggered a series of Ca^2+^-mediated spikes (Fig. 6A). At similar membrane potentials used throughout this study (~-70 mV), conditioning steps that evoked repeated Ca^2+^ spikes triggered persistent firing (spontaneous Ca^2+^ spikes following of the offset of the depolarizing current injection) which were abolished by Terf (Fig. 6B-C). Even with voltage-gated Na^+^ channels blocked with TTX, Terf increased steady-state input resistance (Fig. 6D), consistent with a primary action on ERG channels rather than indirectly affecting excitability via interactions with Na^+^ channels. The effect of Terf on persistent Ca^2+^ spiking did not reflect a diminished stimulus since there was no reduction in the number of Ca^2+^ spikes evoked by the conditioning step (Fig. 6E). In TTX and CCh, depolarizing conditioning steps still evoked a large transient increase in input resistance that was greatly attenuated by Terf (Fig. 6F-G). We found a similar effect of Terf in reducing the transient increase in input resistance assayed with trains of both positive and negative current test pulses (Fig. 6H), arguing that this experiment reflected the underlying neuronal input resistance rather than rectification properties of other currents triggered by the conditioning step (c.f., 62, 63) or artifacts related the detrending procedure we employed.

**Figure 6:**
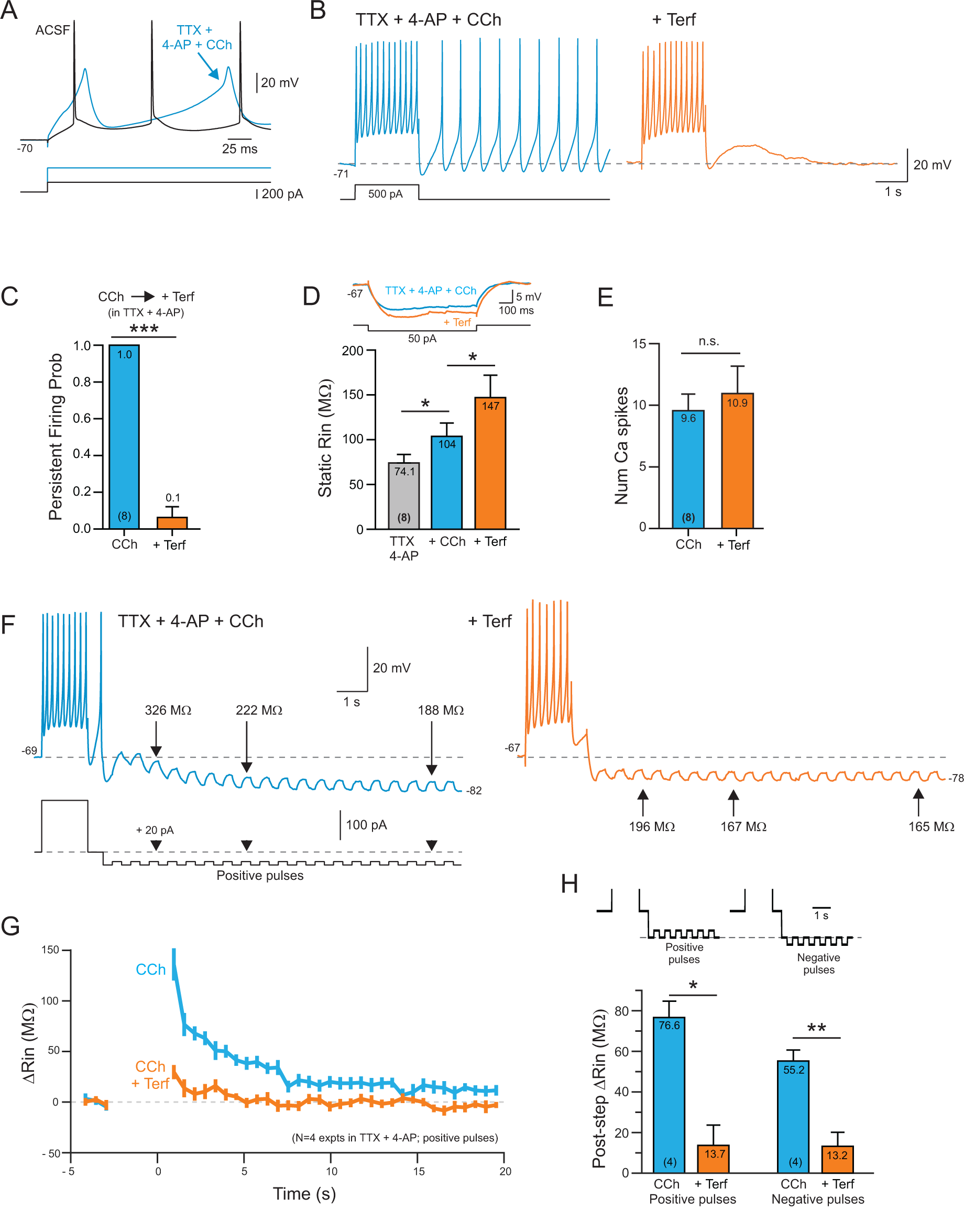
Terfenadine-sensitive persistent firing in the absence of voltage-gated Na+ channels. A, C ± spikes evoked by depolarizing steps following application of 1 *μ*M TTX, 100 *μ*M 4-AP and 2 *μ*M CCh. B, Terfenadine blocks persistent Ca2+ spiking activity triggered by a depolarizing step. C, Plot of the probability of triggering persistent Ca2+ spikes before and after Terf treatment. *** P = 1.41E-06, T = 15, df = 7, paired t-test. D, Plot of input resistance assayed using a single hyperpolarizing step in TTX + 4-AP, following CCh treatment (blue) and following the subsequent addition of Terf (orange). Example step responses shown above plot. * (TTX/CCh) P = 0.0463, T = 2.89, df = 7; * (CCh/CCh+Terf) P = 0.0458, T = 2.90, df = 7; paired t-test. E, Plot of the number of Ca^2+^ spikes evoked by the conditioning step before and after Terf (P = 0.36; paired t-test). F, The underlying ADP response in TTX + 4-AP + CCh is associated with an increase in input resistance assayed using trains of positive current pulses. Terfenadine attenuates both the ADP and the related increase in input resistance. Example R_In_ estimates indicated in panel are detrended. G, Summary plot of change in input resistance from 4 experiments similar to F using positive current test pulses. H, Summary of change in input resistance following conditioning depolarizing step using both trains of positive and negative current pulses. * (positive pulses) P = 0.023, T = 4.31, df = 3; ** (negative pulses) P = 0.015, T = 5.12, df = 3; paired t-test.

Our results suggest that ERG channel blockers are effective in abolishing persistent firing because the conditioning depolarizing step functioned to reduce the steady-state ERG current, leading to a transient hyperexcitable period following the step. This hypothesis would explain the Terf-sensitive increase in input resistance following the conditioning step (Figs. 5B-C and 6F-H) and the enhanced responses to depolarizing test pulses in Fig. 5A. We next sought to test this hypothesis by employing a slow current-clamp ramp protocol (Fig. 7A-E) to reveal how the I/V relationship was altered by the conditioning depolarizing step. In control ACSF, we found essentially no difference in the I/V relationship following the conditioning step (black trace in Fig. 7A shows the subtraction of the ramp response following the conditioning step (“step”) from the response to an identical ramp stimulus presented in isolation; “no step”). In CCh (and without TTX/TEA), the same protocol revealed a negative slope in the I/V relationship in 4/4 experiments. This negative slope response likely reflected a reduction in a leak K^+^ current–rather than an inward current–given its reversal near the K^+^ equilibrium potential. In separate experiments, we applied the GIRK activator ML297 (0.67 *μ*M; without CCh) which generated an outward current in the difference I/V relationship that reversed polarity at approximately the same potential. Results from these experiments are summarized in Figs. 7D-E and are consistent with a transient reduction in ERG current contributing to the hyperexcitability following the conditioning depolarizing step in CCh.

**Figure 7:**
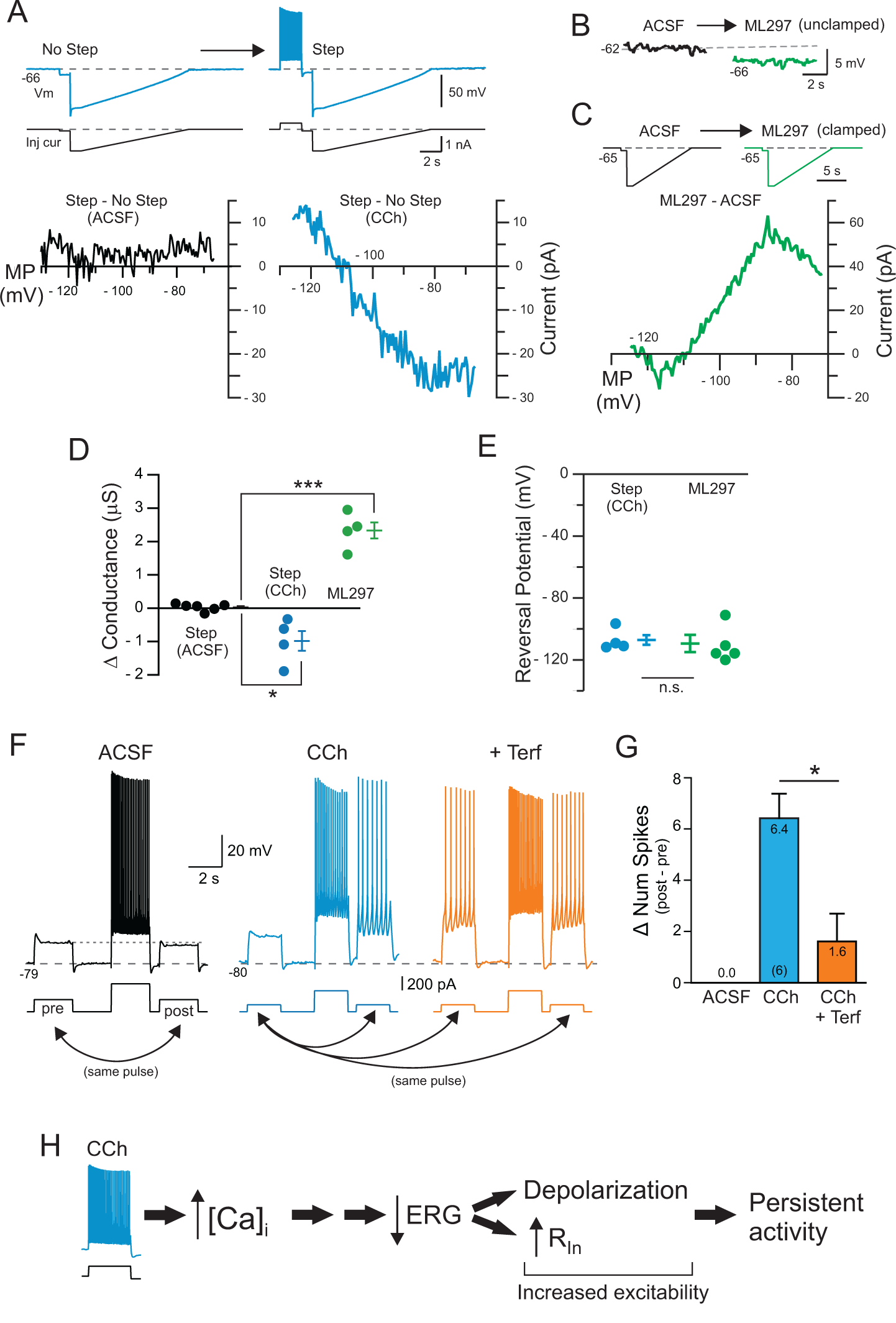
Reduction in leak ERG current contributes to increased excitability following depolarizing step. A, Example responses to ramp current injections with and without a preceding depolarizing current step. Bottom panels show difference I/V responses obtained by subtracting the step + ramp response from the ramp response alone (see Methods for details). Black curve recorded under control (ACSF) conditions and blue curve in CCh. B, Bath application of the GIRK activator ML297 (0.66 *μ*M) hyperpolarizes L5 pyramidal cells. C, Difference I/V response calculated by subtracting ramp response in ML297 from ramp response recorded in ACSF. Initial holding potential adjusted to -65 mV under both conditions. Current injection protocol shown above I/V plot. D, Summary of estimated conductance evoked by depolarizing current steps in control (black symbols), in CCh (blue) and following ML297 application (green). * (step ACSF/CCh) P = 0.0123, T = 3.69, df = 8; *** (ML297) P = 1.45E-05, T = 10.21, df = 8; Two sample t-test. E, Plot of reversal potential of difference I/V plots. n.s. (P = 0.77, two-sample t-test). F, Modulation of responses to weak test depolarizing pulses by conditioning depolarizing step response. Modulation in ACSF assayed in separate experiments than CCh/CCh+Terf. G, Summary plot of change in the number of spikes evoked by two test pulses under each condition (post-step minus pre-step). * P = 0.0199, T = 3.37, df = 5; paired t-test. H, Diagram of potential cascade evoked by the conditioning depolarizing step that leads to reduction in the component of the leak K^+^ current mediated by ERG channels. The loss of part of the standing K^+^ current can account for both the depolarization of pyramidal cells and the transient increase in input resistance.

As another test of the role of ERG modulation in contributing to post-step hyperexcitability, we applied two identical just-subthreshold depolarizing test pulses, one prior to the conditioning step and the other 0.5 s following the conditioning step. In control conditions (ACSF), the response to the second test step (“post”) was slightly diminished compared to the first test step (“pre”; black trace in Fig. 7F). In CCh, the response to the second test step was enhanced and generated a train of APs (blue trace in Fig. 7F). When Terf was applied in combination with CCh, the previously-subthreshold first test step triggered a train of APs which was only slightly increased following the conditioning step (8 spikes on the post step vs 7 spikes in response to the initial test stimulus; orange trace in Fig. 7F). Figure 7G summarizes 6 experiments similar to the one shown in Fig. 7F and suggests that blockade of ERG current with Terf not only increases the steady-state excitability of neocortical pyramidal cells (accounting for the increased response to the first test step) but also occludes the ability of the conditioning depolarizing step to transiently increase intrinsic excitability.

A proposed mechanism for post-conditioning step hyperexcitability is diagrammed in Fig. 7H and postulates that increases in intracellular Ca^2+^ generated in response to the conditioning step leads to reduction the component of leak K^+^ current mediated by ERG channels. The increased post-conditioning step excitability likely reflects both the steady-state depolarization triggered by the reduced leak ERG current as well as the increased input resistance. In most of our experiments we applied a hyperpolarizing bias current soon after the conditioning step to prevent prolonged periods of persistent firing. Without this intervention, the period of post-conditioning step hyperexcitability presumably would last longer as Ca^2+^ accumulations triggered by additional Na^+^ spikes re-engages the same modulatory mechanism, likely leading the persistently attenuated ERG current.

### ERG-mediated hyperexcitability in prefrontal cortical neurons

Finally, we tested whether leak ERG channels contribute to the intrinsic excitability of pyramidal cells in medial prefrontal cortex, where persistent firing is commonly recorded during working memory tasks in both rodents and primates (2, 64, 65, 66). Blockade of ERG current with Terf increased the number of APs evoked by depolarizing steps in L5 mPFC pyramidal cells held at ~-70 mV (Fig. 8A). As with TeA neocortical neurons, Terf increased both the average number of spikes (by 40%; Fig. 8B) and the resting input resistance (by 36%; Fig. 8C) in L5 mPFC pyramidal cells. Terfenadine also abolished persistent firing evoked in 2 μM CCh in 6 of 6 experiments (Fig. 8D-E). Using the same protocol presented Fig. 5B-C, we found a similar increase in input resistance following the conditioning depolarizing step which was greatly reduced by Terf in 4/4 experiments (Fig. 8F-G; results reflecting the same dual R_In_ correction procedure outlined above). The maximal increase in apparent input resistance was somewhat smaller in mPFC than TeA L5 pyramidal cells ~36 vs 25 MΩ, suggesting that ERG-mediated persistent firing may be more robust in TeA than PFC neurons. In both TeA and PFC neurons, Terf abolished persistent firing without affecting either the AP threshold (Fig. 8H) or the AP half-width (Fig. 8I), consistent with a relatively specific action of this agent on a slowly-activating K^+^ current.

**Figure 8:**
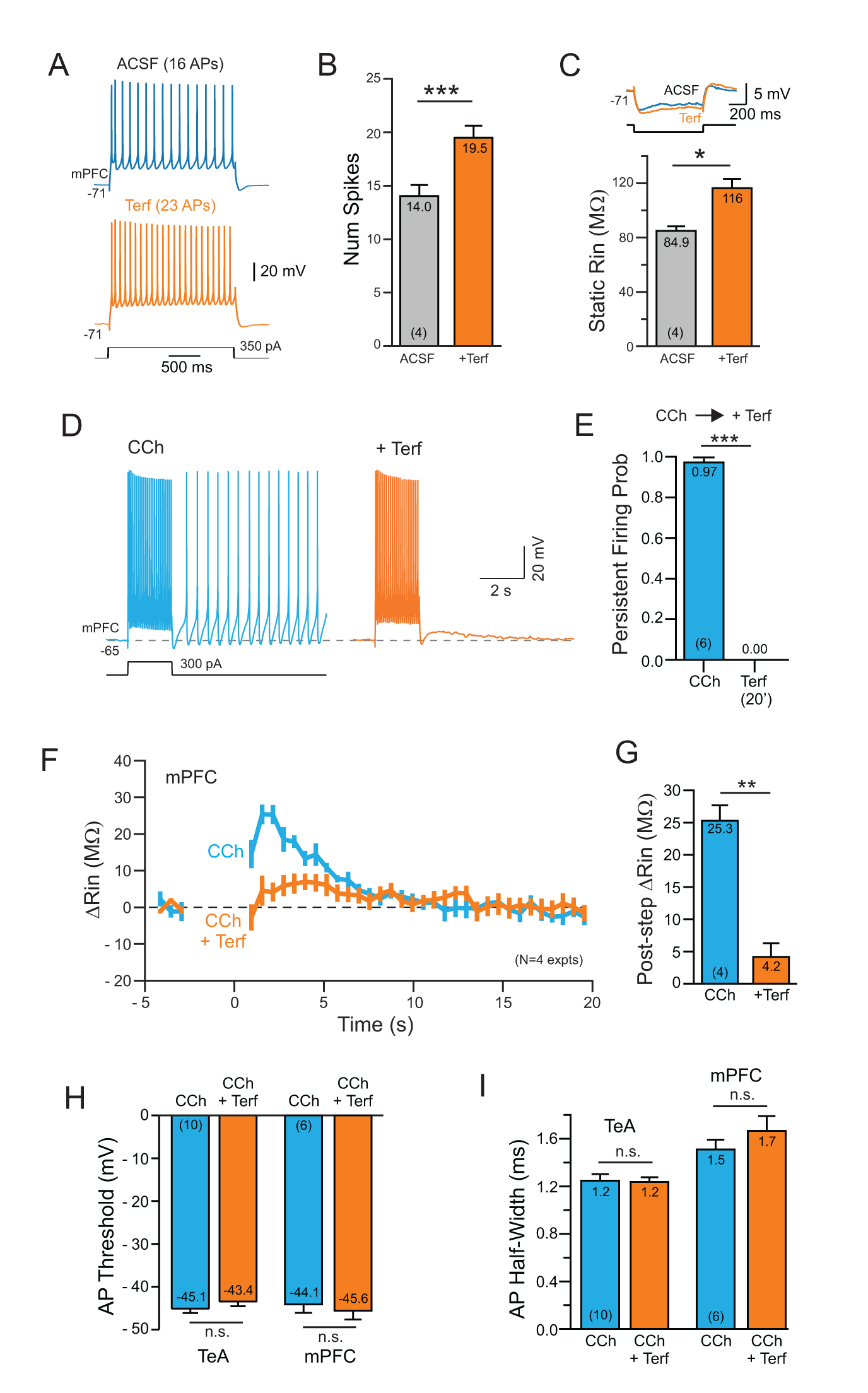
ERG-mediated persistent activity in prefrontal cortical neurons. A, Terfenadine increases the number of APs evoked by depolarizing steps in L5 medial prefrontal pyramidal (mPFC) cells. B, Summary plot of number of APs evoked by the conditioning step in ACSF and Terf. * P = 0.0131, T = 5.30, df = 3, paired t-test. C, Plot of increase in input resistance with Terf. Example responses above plot. *** P = 8.52E-05, T = 29.54, df= 3, paired t-test. D, Terfenadine abolishes persistent firing evoked in CCh. E, Summary plot of effectiveness of Terf in blocking persistent firing in CCh in 6 experiments. *** P = 3.62E-07, T = 34.93, df = 5, paired t-test. F, Plot of the change in input resistance in 4 mPFC pyramidal cells following conditioning depolarizing steps in CCh and CCh+Terf. Experimental protocol the same as shown in Fig. 5B-C. G, Summary plot of maximal increase in input resistance observed during ADP response. ** P = 0.00163, T = 10.95, df = 3; paired t-test. H, Plot of AP threshold in CCh before and after Terf. Left column pair in temporal association (TeA) neocortex and right pair in medial prefrontal cortex. TeA: P = 0.25; PFC: P = 0.40; paired t-test. I, Plot of AP half-width calculated at the half-maximal AP amplitude. Same column order as H. TeA: P = 0.78; PFC: P = 0.33, paired t-test.

## Discussion

We make three principal conclusions in this report, all of which we believe have not been reported previously. First, we find that tonically-active ERG currents contribute to both setting the resting membrane potential and regulating the number of APs triggered in response to depolarizing stimuli. This result, observed in both TeA and PFC pyramidal cells, supports the hypothesis that “leak” ERG K^+^ currents play an important role in shaping the intrinsic physiology cortical neurons. Second, using both biophysical and pharmacological assays, we find that a reduction in leak ERG current appears to play a central role in mediating persistent firing in neocortical neurons. Apparent input resistance increases during the ADP that underlies persistent firing–a result that is inconsistent with the expected decrease in input resistance in I_CAN_-mediated afterdepolarizations (16, 14, 17). Finally, we find that depolarizing stimuli when presented in combination with m1 receptor activation triggers a transient increase in excitability by attenuating leak ERG currents through a Ca^2+^-dependent mechanism. Together, these results point to modulation of leak ERG current as a central underlying mechanism responsible for persistent firing in neocortical neurons and a novel therapeutic target for neurological and psychiatric diseases.

### Potential mechanism of post-stimulus hyperexcitability

Our results suggests that a component of the increased excitability responsible for persistent firing modes in neocortical neurons arises from a rapid attenuation in ERG K^+^ currents that contribute to the leak conductance in pyramidal cells. Following cholinergic receptor activation, depolarizing stimuli appear to attenuate leak ERG current, leading to both membrane depolarizing and an increase in input resistance. Our results provide three principal lines of evidence for ERG contributing to the ADP that underlies persistent firing in neocortical neurons: (1) we directly assayed input resistance during the ADP response and observed a large (30-40%) increase even after compensating for artifacts related to variable R_In_ estimates at different voltages and the ability of subthreshold Na^+^ channels to influence R_In_ measurements, (2) the I/V relationship assayed during the ADP reversed near the K^+^ equilibrium potential and (3) three chemically diverse ERG blockers abolished persistent firing while attenuating both the underlying ADP and the increase in apparent input resistance following the conditioning step.

The ERG-based model of persistent firing is reminiscent of classic studies of the mechanism of cholinergic hyperexcitatibilty (13, 37, 42) which suggested a central role for attenuated leak K^+^ current in promoting persistent firing. This explanation was supplanted by the I_CAN_ model when Alonso and others (e.g., 16, 11) found that the TRP blocker flufenamic acid (FFA) abolished persistent firing while attenuated the underlying ADP in a variety of neurons. Unfortunately, FFA affects many ionic currents, including ERG (67, 68), making it less useful in discriminating between potential mechanisms. Another commonly used TRP channel blocker, SKF-96365, blocks heterologously-expressed ERG channels; 69.

Our results from assays of input resistance and the the I/V properties of the ADP response are inconsistent with recent ICAN-based models of persistent activity (e.g., 40, 17, 11, 19, 18) which predict that persistent firing should be accompanied by a decrease (rather than an increase) in input resistance during the ADP. However, several previous investigators observed increases in apparent input resistance during the ADP (14) which they attributed to rectification properties of TRP channels (62, 63). This prior result provided the motivation for our two-step R_In_ correction procedure and for assays of input resistance using both positive- and negative-going current steps that were feasible once voltage-gated Na^+^ channels were blocked, as in the experiments in Fig. 6. Andrade and colleagues (14) observed that Cs^+^ ions failed to abolish persistent activity. While we confirmed this result, this finding is difficult to interpret since ERG channels–unlike most voltage-gated K^+^ channels–are not blocked by Cs^+^ (70). Nevertheless, without further experiments we cannot exclude that subthreshold Na^+^ channel and I_CAN_ mechanisms contribute to persistent firing, along with ERG, since we consistently find a residual small ADP following treatment with ERG blockers. However, our results suggest that in 2 *μ*M CCh, attenuating ERG using Terf is sufficient to trigger persistent firing from -70 mV. Based on previous studies using heterologously-expressed ERG channels, we expect that our pharmacological treatments blocked 50-80% of whole-cell ERG current (e.g., IC_50_ for ErgToxin1 is 10-60 nM; 45, 52). Until new molecular or pharmacological tools become available, we cannot determine whether the residual ADP represented unblocked ERG current or contributions from other mechanisms.

While our results suggest that attenuation of leak ERG current appears to be a critical component in intrinsic persistent spiking responses, we did not attempt to define the specific biophysical mechanisms responsible for ERG modulation. In other systems, ERG currents are inhibited both by protein kinase C (PKC)-mediated phosphorylation (24) and by depleting phosphatidylinositol 4,5-bisphosphate (PIP_2_; 71). Activation of muscarinic m1 receptors activates phospholipase C (PLC) which hydrolyzes PIP_2_ into diacylglycerol (DAG) and inositol-1,4,5-triphosphate (IP_3_), potentially enabling both PIP_2_ and PKC mechanisms. Understanding how the transient elevation of intracellular Ca^2+^ concentration during the conditioning step leads to a further decrease in ERG current is a central but difficult question to resolve. The rapid onset of persistent firing (and the underlying ADP) would suggest a dominant role of PIP_2_-mediated modulation since FRET-based measurements suggest that that process can operate within ~1 s (72). It is possible that PIP_2_ depletion leads to the initial post-conditioning step increase in excitability which is then reinforced by subsequent PKC-mediated phosphorylation of ERG channels. Alternatively, the normally slow actions of protein kinases could be accelerated if they were “primed” via scaffolding proteins such as caveolin (73, 74). Human ERG channels can interact with caveolin (75, 76), providing a potentially rapid pathway for G-protein stimulated phosphorylation. It is also possible that the modulation of ERG affects primarily gating properties rather than reducing channel conductance, as suggested by previous work (71, 54, 24). Discriminating between these potential mechanisms, and revealing how Ca^2+^ transients function to trigger persistent firing when combined with muscarinic receptor activation, will likely require development of new rapid FRET-based tools that are beyond the scope of the present study.

Molecular genetic tests of the role of ERG channels in neocortical neurons will likely require conditional knockouts of one or more ERG genes as permanent deletion of these proteins is lethal in mice at E11.5 (77, 78). To our knowledge, conditional ERG knockout mice have not been generated. While more extensive voltage-clamp analysis would help clarify the functional role of ERG channels, those experiments are challenging to undertake in CNS neurons since ERG channel properties are affected by both changes in Mg^2+^ and K^+^ concentration (79, 80, 57, 81). ERG channels are also a frequent nonspecific target of pharmacological agents, including many antipsychotics (32). ERG channels are permeable to Cs^+^ ions (70), further complicating conventional voltage-clamp based current analysis.

### Functional significance of ERG-mediated intrinsic persistent firing

While the origin of persistent activity associated with short-term memory and other cognitive functions (2, 3, 82) has not been determined, the identification of ERG modulation as a potential intrinsic mechanism should facilitate determining of the role of intrinsic vs recurrent network mechanisms. The availability of highly specific ERG channel blockers (e.g., ErgToxin1) makes it feasible to directly test to what extent intrinsic biophysical properties contribute to commonly-observed persistent firing modes such as delay period firing working memory tasks. ERG-blocking agents also could be used to determine whether persistently-active subcortical circuits (e.g., the VOR) rely on modulation of ERG current.

Cholinergic receptor stimulation has long been known to increase neuronal excitability and promote intrinsic persistent firing modes (e.g., 13). Since cholinergic agents strongly influence cognitive processes associated with persistent firing, such as working memory (2), a fundamental question arising from previous in vitro studies is how intrinsic and circuit mechanisms could be integrated to generate stimulus-specific persistent firing. Modulating most types of intrinsic currents that contribute to the resting (leak) conductance of neurons would be expected to dramatically alter tuning of synaptic weights within recurrent networks (82). Because of its slow activation kinetics (54, 24), ERG-mediated persistence is an attractive intrinsic mechanism to co-exist with precisely-tuned synaptic networks. Presumably brief synaptically-evoked discharges will be only weakly affected by ERG currents– leaving delicate network tuning unaffected–while stronger, or more sustained, synaptic excitation would be preferentially amplified by intrinsic currents. Through this mechanism, ERG-mediated intrinsic persistent activity could function to “tag” the most active subset of of cells within a larger neuronal ensemble driven by a stimulus. A central predication of this hypothesis is that ERG blockers should preferentially attenuate late phases of neuronal discharges as well as reducing post-stimulus persistent activity.

Recent genetic studies have suggested an association between single nucleotide polymorphism (SNP) in ERG channels and schizophrenia (31; 83). The mutated ERG channel, Kv11.1-3.1, has altered gating properties comparing to its wild type counterpart (84) and is highly expressed in cortical neurons within a subpopulation of schizophrenic patients (30). Since schizophrenia is often associated with altered PFC activity and impaired working memory function (85), it is possible that a component of this disease reflects abnormal ERG function which could result in changes in both the average discharge rate of cortical neurons and the regularity of their firing. Many common second-generation antipsychotic mediations, such as risperidone, are potent ERG blockers (32) and patients with ERG mutations appear to be preferentially responsive to ERG-blocking antipsychotics (86). Since our study suggests that ERG is an important component of the normal constellation of “leak” K^+^ channels in at least a subset of neocortical neurons (regular spiking deep pyramidal cells), our results provide a novel cellular mechanism for the actions of many second-generation antipsychotics that could help explain cognitive dysfunction associated with schizophrenia.

## Acknowledgments

We thank Todd Pressler and Hannah Arnson and for helpful discussions and Chris Ford, Rodrigo Andrade and Diana Kunze for providing constructive comments on this study. This work was supported by NIH grant R01-DC04285 to B.W.S.

This PDF document (v1.11) was generated using Pandoc on June 26, 2017.

## References

1. Major, G., Baker, R., Aksay, E., Seung, H. S. & Tank, D. W. Plasticity and tuning of the time course of analog persistent firing in a neural integrator. Proceedings of the National Academy of Sciences of the United States of America 101, 7745–50 (2004).

2. Fuster, J. M. & Alexander, G. E. Neuron Activity Related to Short-Term Memory. 173, 652–654 (1971).

3. Eichenbaum, H. Time cells in the hippocampus: a new dimension for mapping memories. Nature reviews. Neuroscience 15, 732–44 (2014).

4. Hyde, R. A. & Strowbridge, B. W. Mnemonic representations of transient stimuli and temporal sequences in the rodent hippocampus in vitro. Nature neuroscience 15, 1430–8 (2012).

5. Larimer, P. & Strowbridge, B. W. Representing information in cell assemblies: persistent activity mediated by semilunar granule cells. Nature Neuroscience 13, 213–222 (2010).

6. Wang, X. J. Synaptic basis of cortical persistent activity: the importance of NMDA receptors to working memory. The Journal of neuroscience: the official journal of the Society for Neuroscience 19, 9587–9603 (1999).

7. Hebb, D. The organization of behavior: a neuropsychological theory. (John Wiley & Sons, 1949).

8. Hopfield, J. J. Neural networks and physical systems with emergent collective computational abilities. Proceedings of the National Academy of Sciences of the United States of America 79, 2554–2558 (1982).

9. Andrade, R. Cell excitation enhances muscarinic cholinergic responses in rat association cortex. Brain research 548, 81–93 (1991).

10. Haj-Dahmane, S. & Andrade, R. Muscarinic activation of a voltage-dependent cation nonselective current in rat association cortex. The Journal of neuroscience: the official journal of the Society for Neuroscience 16, 3848–3861 (1996).

11. Rahman, J. & Berger, T. Persistent activity in layer 5 pyramidal neurons following cholinergic activation of mouse primary cortices. European Journal of Neuroscience 34, 22–30 (2011).

12. Pressler, R. T., Inoue, T. & Strowbridge, B. W. Muscarinic receptor activation modulates granule cell excitability and potentiates inhibition onto mitral cells in the rat olfactory bulb. The Journal of neuroscience: the official journal of the Society for Neuroscience 27, 10969–81 (2007).

13. Krnjevic, K., Pumain, R. & Renaudt, L. The mechanism of excitation by acetylcholine in the cerebral cortex. J Physiol 215, 247–268 (1971).

14. Haj-Dahmane, S. & Andrade, R. Ionic mechanism of the slow afterdepolarization induced by muscarinic receptor activation in rat prefrontal cortex. Journal of neurophysiology 80, 1197–1210 (1998).

15. Richardson, R. T. & DeLong, M. R. Nucleus basalis of Meynert neuronal activity during a delayed response task in monkey. Brain Research 399, 364–368 (1986).

16. Egorov, A. V., Hamam, B. N., Fransen, E., Hasselmo, M. E. & Alonso, A. A. Graded Persistent activity in entorhinal cortex neurons. Nature 420, 173–178 (2002).

17. Fransén, E., Tahvildari, B., Egorov, A. V., Hasselmo, M. E. & Alonso, A. A. Mechanism of Graded Persistent Cellular Activity of Entorhinal Cortex Layer V Neurons. Neuron 49, 735–746 (2006).

18. Jochems, A. & Yoshida, M. Persistent firing supported by an intrinsic cellular mechanism in hippocampal CA3 pyramidal cells. European Journal of Neuroscience 38, 2250–2259 (2013).

19. Zhang, Z., Reboreda, A., Alonso, A., Barker, P. A. & Séguéla, P. TRPC channels underlie cholinergic plateau potentials and persistent activity in entorhinal cortex. Hippocampus 21, 386–397 (2011).

20. Nilius, B. & Owsianik, G. The transient receptor potential family of ion channels. Genome Biol 12, 218 (2011).

21. Yamada-Hanff, J. & Bean, B. P. Persistent Sodium Current Drives Conditional Pacemaking in CA1 Pyramidal Neurons under Muscarinic Stimulation. Journal of Neuroscience 33, 15011–15021 (2013).

22. Delmas, P. & Brown, D. a. Pathways modulating neural KCNQ/M (Kv7) potassium channels. Nature reviews. Neuroscience 6, 850–862 (2005).

23. Schnee, M. E. & Brown, B. S. Selectivity of linopirdine (DuP 996), a neurotransmitter release enhancer, in blocking voltage-dependent and calcium-activated potassium currents in hippocampal neurons. J Pharmacol Exp Ther 286, 709–717 (1998).

24. Cockerill, S. L., Tobina, B., Torrecilla, I., Willars, G. B., Standen, N. B. & Mitcheson, J. S. Modulation of hERG potassium currents in HEK-293 cells by protein kinase C. Evidence for direct phosphorylation of pore forming subunits. The Journal of physiology 581, 479–493 (2007).

25. Papa, M., Boscia, F., Canitano, A., Castaldo, P., Sellitti, S., Annunziato, L. & Taglialatela, M. Expression pattern of the ether-a-gogo-related (ERG) K+ channel-encoding genes ERG1, ERG2, and ERG3 in the adult rat central nervous system. Journal of Comparative Neurology 466, 119–135 (2003).

26. Saganich, M. J., Machado, E. & Rudy, B. Differential Expression of Genes Encoding Subthreshold-Operating Voltage-Gated K+ Channels in Brain. The Journal of Neuroscience 21, 4609–4624 (2001).

27. Ji, H., Tucker, K. R., Putzier, I., Huertas, M. A., Horn, J. P., Canavier, C. C., Levitan, E. S. & Shepard, P. D. Functional characterization of ether-a-go-go-related gene potassium channels in midbrain dopamine neurons - implications for a role in depolarization block. European Journal of Neuroscience 36, 2906–2916 (2012).

28. Hardman, R. M. & Forsythe, I. D. Ether-à-go-go-related gene K+ channels contribute to threshold excitability of mouse auditory brainstem neurons. The Journal of physiology 587, 2487–97 (2009).

29. Fano, S., Çalişkan, G. & Heinemann, U. Differential effects of blockade of ERG channels on gamma oscillations and excitability in rat hippocampal slices. European Journal of Neuroscience 36, 3628–3635 (2012).

30. Huffaker, S. J., Chen, J., Nicodemus, K. K., Sambataro, F., Yang, F., Mattay, V., Lipska, B. K., Hyde, T. M., Song, J., Rujescu, D., Giegling, I., Mayilyan, K., Proust, M. J., Soghoyan, A., Caforio, G., Callicott, J. H., Bertolino, A., Meyer-Lindenberg, A., Chang, J., Ji, Y., Egan, M. F., Goldberg, T. E., Kleinman, J. E., Lu, B. & Weinberger, D. R. A primate-specific, brain isoform of KCNH2 affects cortical physiology, cognition, neuronal repolarization and risk of schizophrenia. Nature medicine 15, 509–18 (2009).

31. Atalar, F., Acuner, T. T., Cine, N., Oncu, F., Yesilbursa, D., Ozbek, U. & Turkcan, S. Two four-marker haplotypes on 7q36.1 region indicate that the potassium channel gene HERG1 (KCNH2, Kv11.1) is related to schizophrenia: a case control study. Behavioral and brain functions: BBF 6, 27 (2010).

32. Wible, B. A., Hawryluk, P., Ficker, E., Kuryshev, Y. A., Kirsch, G. & Brown, A. M. HERG-Lite(R): A novel comprehensive high-throughput screen for drug-induced hERG risk. Journal of Pharmacological and Toxicological Methods 52, 136–145 (2005).

33. Connors, B. W. & Gutnick, M. J. Intrinsic firing patterns of diverse neocortical neurons. Trends in neurosciences 13, 99–104 (1990).

34. Dégenètais, E., Thierry, A.-M., Glowinski, J. & Gioanni, Y. Electrophysiological properties of pyramidal neurons in the rat prefrontal cortex: an in vivo intracellular recording study. Cerebral cortex (New York, N.Y.: 1991) 12, 1–16 (2002).

35. Schubert, D., Staiger, J. F., Cho, N., Kötter, R., Zilles, K. & Luhmann, H. J. Layer-specific intracolumnar and transcolumnar functional connectivity of layer V pyramidal cells in rat barrel cortex. The Journal of neuroscience: the official journal of the Society for Neuroscience 21, 3580–3592 (2001).

36. Yoshida, M. & Hasselmo, M. E. Persistent Firing Supported by an Intrinsic Cellular Mechanism in a Component of the Head Direction System. Journal of Neuroscience 29, 4945–4952 (2009).

37. McCormick, D. A. & Prince, D. A. Two types of muscarinic response to acetylcholine in mammalian cortical neurons. Proceedings of the National Academy of Sciences of the United States of America 82, 6344–6348 (1985).

38. Gulledge, A. T., Bucci, D. J., Zhang, S. S., Matsui, M. & Yeh, H. H. M1 receptors mediate cholinergic modulation of excitability in neocortical pyramidal neurons. J. Neurosci. 29, 9888–9902 (2009).

39. Winograd, M., Destexhe, A. & Sanchez-Vives, M. V. Hyperpolarization-activated graded persistent activity in the prefrontal cortex. Proceedings of the National Academy of Sciences of the United States of America 105, 7298–7303 (2008).

40. Knauer, B., Jochems, A., Valero-Aracama, M. J. & Yoshida, M. Long-lasting intrinsic persistent firing in rat CA1 pyramidal cells: A possible mechanism for active maintenance of memory. Hippocampus 23, 820–831 (2013).

41. Greene, C. C., Schwindt, P. C. & Crill, W. E. Properties and Ionic Mechanisms of a Metabotropic Glutamate Receptor-Mediated Slow Afiierdepolarization in Neocortical Neurons. 72, 693–704 (1994).

42. McCormick, D. A. & Prince, D. A. Mechanisms of action of acetylcholine in the guinea-pig cerebral cortex in vitro. J. Physiol. (Lond.) 375, 169–194 (1986).

43. Shepard, P. D., Canavier, C. C. & Levitan, E. S. Ether-a-go-go-related gene potassium channels: What’s all the buzz about? Schizophrenia Bulletin 33, 1263–1269 (2007).

44. Sacco, T., Bruno, A., Wanke, E. & Tempia, F. Functional roles of an ERG current isolated in cerebellar Purkinje neurons. Journal of neurophysiology 90, 1817–1828 (2003).

45. Hill, A. P., Sunde, M., Campbell, T. J. & Vandenberg, J. I. Mechanism of block of the hERG K+ channel by the scorpion toxin CnErg1. Biophysical journal 92, 3915–29 (2007).

46. Scherer, C. R., Lerche, C., Decher, N., Dennis, A. T., Maier, P., Ficker, E., Busch, A. E., Wollnik, B. & Steinmeyer, K. The antihistamine fexofenadine does not affect I(Kr) currents in a case report of drug-induced cardiac arrhythmia. British journal of pharmacology 137, 892–900 (2002).

47. Kamiya, K., Niwa, R., Mitcheson, J. S. & Sanguinetti, M. C. Molecular Determinants of hERG Channel Block. 69, 1709–1716 (2006).

48. Spector, P. S., Curran, M. E., Keating, M. T. & Sanguinetti, M. C. Class III antiarrhythmic drugs block HERG, a human cardiac delayed rectifier K+ channel. Open-channel block by methanesulfonanilides. Circulation research (1996).

49. Chtcheglova, L. A., Atalar, F., Ozbek, U., Wildling, L., Ebner, A. & Hinterdorfer, P. Localization of the ergtoxin-1 receptors on the voltage sensing domain of hERG K+ channel by AFM recognition imaging. Pflugers Archiv European Journal of Physiology 456, 247–254 (2008).

50. Milnes, J. T., Dempsey, C. E., Ridley, J. M., Crociani, O., Arcangeli, A., Hancox, J. C. & Witchel, H. J. Preferential closed channel blockade of HERG potassium currents by chemically synthesised BeKm-1 scorpion toxin. FEBS Letters 547, 20–26 (2003).

51. Vandenberg, J. I., Torres, Æ. A. M., Campbell, T. J. & Kuchel, Æ. P. W. The HERG K + channel: progress in understanding the molecular basis of its unusual gating kinetics. 89–97 (2004). doi:10.1007/s00249-003-0338-3

52. Pardo-López, L., García-Valdés, J., Gurrola, G. B., Robertson, G. A. & Possani, L. D. Mapping the receptor site for ergtoxin, a specific blocker of ERG channels. FEBS Letters 510, 45–49 (2002).

53. Niculescu, D., Hirdes, W., Hornig, S., Pongs, O. & Schwarz, J. R. Erg potassium currents of neonatal mouse Purkinje cells exhibit fast gating kinetics and are inhibited by mGluR1 activation. J. Neurosci. 33, 16729–16740 (2013).

54. Hirdes, W., Horowitz, L. F. & Hille, B. Muscarinic modulation of erg potassium current. The Journal of physiology 559, 67–84 (2004).

55. Lin, M. C. A. & Papazian, D. M. Differences between ion binding to eag and herg voltage sensors contribute to differential regulation of activation and deactivation gating. Channels 1, 429–437 (2007).

56. Prole, D. L., Lima, P. a & Marrion, N. V. Mechanisms underlying modulation of neuronal KCNQ2/KCNQ3 potassium channels by extracellular protons. The Journal of general physiology 122, 775–793 (2003).

57. Wang, S., Liu, S., Morales, M. J., Strauss, H. C. & Rasmusson, R. L. A quantitative analysis of the activation and inactivation kinetics of HERG expressed in Xenopus oocytes. J Physiol (Lond) 502, 45–60 (1997).

58. Zhou, Z., Gong, Q., Ye, B., Fan, Z., Makielski, J. C., Robertson, G. a & January, C. T. Properties of HERG channels stably expressed in HEK 293 cells studied at physiological temperature. Biophysical journal 74, 230–41 (1998).

59. Robinson, R. B. & Siegelbaum, S. A. Hyperpolarization-activated cation currents: from molecules to physiological function. Annual Review of Physiology 65, 453–480 (2003).

60. Notomi, T. & Shigemoto, R. Immunohistochemical Localization of Ih Channel Subunits, HCN1-4, in the Rat Brain. Journal of Comparative Neurology 471, 241–276 (2004).

61. Crill, W. E. W. Persistent sodium current in mammalian central neurons. Annual Review of Physiology 58, 349–62 (1996).

62. Inoue, R. & Isenberg, G. Intracellular Calcium Ions Modulate Acetylcholine-induced Inward Current in Guinea-pig Ileum. Journal of Physiology 424, 73–92 (1990).

63. Inoue, R. & Isenberg, G. Effect of membrane potential on acetylcholine-induced inward current in guinea-pig ileum. The Journal of physiology 424, 57–71 (1990).

64. Miyashita, Y. & Chang, H. S. Neuronal correlate of pictorial short-term memory in the primate temporal cortex. Nature 331, 68–70 (1988).

65. Young, B. J., Otto, T., Fox, G. D. & Eichenbaum, H. Memory representation within the parahippocampal region. The Journal of neuroscience: the official journal of the Society for Neuroscience 17, 5183–5195 (1997).

66. Liu, D., Gu, X., Zhu, J., Zhang, X., Han, Z., Yan, W., Cheng, Q., Hao, J., Fan, H., Hou, R., Chen, Z., Chen, Y. & Li, C. T. Medial prefrontal activity during delay period contributes to learning of a working memory task. Science 346, 458–463 (2014).

67. Guinamard, R., Simard, C. & Del Negro, C. Flufenamic acid as an ion channel modulator. Pharmacology and Therapeutics 138, 272–284 (2013).

68. Malykhina, A. P., Shoeb, F. & Akbarali, H. I. Fenamate-induced enhancement of heterologously expressed HERG currents in Xenopus oocytes. Eur. J. Pharmacol. 452, 269–277 (2002).

69. Liu, H., Yang, L., Chen, K. H., Sun, H. Y., Jin, M. W., Xiao, G. S., Wang, Y. & Li, G. R. SKF-96365 blocks human ether-a-go-go-related gene potassium channels stably expressed in HEK 293 cells. Pharmacological Research 104, 61–69 (2016).

70. Zhang, S., Kehl, S. J. & Fedida, D. Modulation of human ether-à-go-go-related K+ (HERG) channel inactivation by Cs+ and K+. The Journal of physiology 548, 691–702 (2003).

71. Bian, J., Cui, J. & McDonald, T. V. HERG K+ Channel Activity Is Regulated by Changes in Phosphatidyl Inositol 4,5-Bisphosphate. Circulation Research 89, 1168–1176 (2001).

72. Jensen, J. B., Lyssand, J. S., Hague, C. & Hille, B. Fluorescence changes reveal kinetic steps of muscarinic receptor-mediated modulation of phosphoinositides and Kv7.2/7.3 K+ channels. J. Gen. Physiol. 133, 347–359 (2009).

73. Isshiki, M. & Anderson, R. G. Calcium signal transduction from caveolae. Cell Calcium 26, 201–208 (1999).

74. Gallego, M., Alday, A., Alonso, H. & Casis, O. Adrenergic regulation of cardiac ionic channels: role of membrane microdomains in the regulation of kv4 channels. Biochim. Biophys. Acta 1838, 692–699 (2014).

75. Ma, Q., Yu, H., Lin, J., Sun, Y., Shen, X. & Ren, L. Screening for cardiac HERG potassium channel interacting proteins using the yeast two-hybrid technique. Cell Biol. Int. 38, 239–245 (2014).

76. Lin, J., Lin, S., Choy, P. C., Shen, X., Deng, C., Kuang, S., Wu, J. & Xu, W. The regulation of the cardiac potassium channel (HERG) by caveolin-1. Biochem. Cell Biol. 86, 405–415 (2008).

77. Teng, G. Q., Zhao, X., Lees-Miller, J. P., Quinn, F. R., Li, P., Rancourt, D. E., London, B., Cross, J. C. & Duff, H. J. Homozygous missense N629D hERG (KCNH2) potassium channel mutation causes developmental defects in the right ventricle and its outflow tract and embryonic lethality. Circulation Research 103, 1483–1491 (2008).

78. Vijayaraj, P., Le Bras, A., Mitchell, N., Kondo, M., Juliao, S., Wasserman, M., Beeler, D., Spokes, K., Aird, W. C., Baldwin, H. S. & Oettgen, P. Erg is a crucial regulator of endocardial-mesenchymal transformation during cardiac valve morphogenesis. Development (Cambridge, England) 139, 3973–85 (2012).

79. Po, S. S., Wang, D. W., Yang, I. C., Johnson, J. P., Nie, L. & Bennett, P. B. Modulation of HERG potassium channels by extracellular magnesium and quinidine. Journal of cardiovascular pharmacology 33, 181–5 (1999).

80. Ho, W. K., Kim, I., Lee, C. O. & Earm, Y. E. Voltage-dependent blockade of HERG channels expressed in Xenopus oocytes by external Ca2+ and Mg2+. Journal of Physiology 507, 631–638 (1998).

81. Sanguinetti, M. C., Jiang, C., Curran, M. E. & Keating, M. T. A mechanistic link between an inherited and an acquired cardiac arrhythmia: hERG encodes the IKr potassium channel. Cell 81, 299–307 (1995).

82. Zylberberg, J. & Strowbridge, B. W. Mechanisms of Persistent Activity in Cortical Circuits: Possible Neural Substrates for Working Memory. Annual Reviews Neuroscience In press (2017).

83. Hashimoto, R., Ohi, K., Yasuda, Y., Fukumoto, M., Yamamori, H., Kamino, K., Morihara, T., Iwase, M., Kazui, H. & Takeda, M. The KCNH2 gene is associated with neurocognition and the risk of schizophrenia. The world journal of biological psychiatry: the official journal of the World Federation of Societies of Biological Psychiatry 14, 114–20 (2013).

84. Heide, J., Mann, S. A. & Vandenberg, J. I. The Schizophrenia-Associated Kv11.1-3.1 Isoform Results in Reduced Current Accumulation during Repetitive Brief Depolarizations. PLoS ONE 7, (2012).

85. Weinberger, D. R. & Berman, K. F. Prefrontal function in schizophrenia: confounds and controversies. Philosophical Transactions of the royal Soceity: Biological Sciences 351, 1495–1503 (1996).

86. Apud, J. A., Zhang, F., Decot, H., Bigos, K. L. & Weinberger, D. R. Genetic variation in KCNH2 associated with expression in the brain of a unique hERG isoform modulates treatment response in patients with schizophrenia. American Journal of Psychiatry 169, 725–734 (2012).

